# Chronic unpredictable stress induces negative cognitive bias in rats with sex-specific changes in neurogenesis and functional connectivity

**DOI:** 10.64898/2026.10.29.762765

**Authors:** Amanda B. Namchuk, Travis E. Hodges, Andrew J. McGovern, Liisa A.M. Galea

## Abstract

**Background:** Females are more likely to be diagnosed with major depressive disorder (MDD) than males. Cognitive symptoms of MDD include negative cognitive bias, the perception of ambiguous situations as negative, which predicts MDD episode onset and severity. MDD is associated with changes to functional connectivity between regions involved in cognition, emotion, attention, and memory.

**Methods:** Here we examined the impact of chronic unpredictable stress (CUS) on negative cognitive bias in male (n=18) and female (n=22) Sprague-Dawley rats. We assessed hippocampal neurogenesis and c-Fos expression in response to an ambiguous context across 15 regions of the frontal cortex, amygdala, nucleus accumbens, and hippocampus, including in newborn dentate gyrus (DG) neurons, to examine functional connectivity during cognitive bias testing. We conducted a graph-based network analysis and constructed linear models to predict freezing in the ambiguous context based on regional activation.

**Results:** CUS-exposed rats froze more to the ambiguous context than non-CUS-exposed rats, regardless of sex. CUS decreased neurogenesis in the ventral DG of both sexes, and increased activation of new neurons in the dorsal DG of males only. In response to the ambiguous context, CUS males had an increase in c-Fos expression in the amygdala. CUS produced extensive, sex-specific network reorganization, decreasing global network efficiency in males but not in females. Freezing in the ambiguous context was predicted by activation of new neurons in combination with the ventral CA1 in opposing directions across sex.

**Conclusions:** These findings suggest that sex-specific pathways drive the expression of CUS-induced negative cognitive bias, underscoring the necessity of sex-stratified preclinical and clinical research on negative cognitive bias in depression.

## INTRODUCTION

Major depressive disorder (MDD) is one of the leading causes of disability worldwide (1). Females are twice as likely to develop MDD compared to males (2), and the severity of multiple MDD symptoms, including rumination (repeatedly focusing on negative thoughts), is greater in females than males (3–5). Negative cognitive bias, the perception of ambiguous stimuli as negative, is associated with more rumination (6) and predicts the onset, maintenance, and recurrence of MDD (7–10), but is resistant to treatment (11–13). However, few studies have examined how sex may influence negative cognitive bias.

Extensive sex differences in brain transcriptomic profiles (14–17) and functional connectivity (18–20) in individuals with MDD have been established. The neural mechanisms underpinning negative cognitive bias in MDD include hyperactivity of limbic regions such as the amygdala and hippocampus and abnormal activity across subregions of the medial and lateral prefrontal cortex (PFC) involved in cognitive control and appraisal of emotional information (21–26), although potential sex differences in this model are unknown. Furthermore, hyperconnectivity between the limbic system (hippocampus, medial PFC, amygdala) and reward regions (nucleus accumbens) predicted negative cognitive bias in a combined-sex sample of individuals with MDD (27). However, to our knowledge, no studies have examined sex differences in the brain correlates of negative cognitive bias. Therefore, this study was designed to examine possible sex differences in the functional connectivity underpinning negative cognitive bias.

Hippocampal integrity is compromised in MDD, including reduced neurogenesis, reduced volume, and altered signaling patterns (28–31). Cognitive bias performance relies on pattern separation (32,33), which requires young adult-born neurons in the dentate gyrus (DG) (34–38). Sex differences in hippocampal neurogenesis are noted throughout adulthood, including after stress exposure and in response to pattern separation tasks (39–41). However, the sex-specific role of hippocampal neurogenesis in negative cognitive bias remains unknown.

Depressive-like endophenotypes, including negative cognitive bias, can be modeled in rodents using exposure to chronic unpredictable stress (32,33,42–47). The majority of preclinical investigations of negative cognitive bias have been performed in males only, but one study reported increased negative cognitive bias after chronic restraint stress in both sexes (32). Here we aim to probe the neural mechanisms underlying chronic unpredictable stress (CUS)-induced negative cognitive bias in both sexes to shed light on this symptom of MDD.

In the current study, we examined cognitive bias in male and female rats after CUS compared to non-CUS-exposed (NC) controls using a fear-based cognitive bias task (48,49). We examined neural activity (c-Fos expression) across 15 regions and in newborn DG neurons in response to the ambiguous context of the cognitive bias task. Graph theory was applied to the c-Fos data to extract network metrics describing the functional networks recruited by each group during cognitive bias testing. Linear modeling identified group differences in the regions in which neural activity best predicted freezing in the ambiguous context. We hypothesized that CUS would increase negative cognitive bias and that distinct patterns of functional connectivity would emerge by sex and stress to predict freezing in the cognitive bias task.

## METHODS AND MATERIALS

### Animals

Male and female Sprague-Dawley rats (N=40) were bred in-house from animals obtained from Charles River (Québec, Canada). Males (n=18) and females (n=22) were group housed (2-3 per cage) in separate colony rooms under a 12h light-dark cycle (lights on 07:00h). Rats were housed in opaque polyurethane bins (48 x 27 x 20 cm) with aspen chip bedding and ad libitum access to autoclaved tap water and rat chow (Jamieson’s Pet Food Distributors Ltd, Delta, BC, Canada). Rats were left undisturbed, apart from weekly cage changing, until they reached the correct age for testing. All experimental procedures were approved by the University of British Columbia Animal Care Committee and in accordance with the Canadian Council on Animal Care guidelines.

### Chronic unpredictable stress

Rats were randomly assigned to either undergo a CUS procedure (adapted from Wainwright et al., 2011) or remain as non-CUS exposed (NC) controls (9 males and 11 females per group). Briefly, starting on postnatal day (PND) 85, CUS animals were exposed to two different stressors pseudorandomly each day for 14 days (Supplemental Table 1). NC rats were housed separately, and remained undisturbed except for weekly cage changing and weighing until beginning the cognitive bias procedure on PND100.

### Cognitive Bias Task Procedure & Tissue Collection

The cognitive bias procedure was performed as previously described (48,49). Briefly, starting 2 days after CUS or NC, rats were placed in a foot shock-paired context (Context A) and in a no-shock-paired context (Context B) for 5 minutes each, daily for 16 days. On Day 18, rats were placed in an ambiguous context partially resembling both Contexts A and B for a 5 min test with no foot shock.

Percent time freezing (no head or body movement besides breathing) during the first 3 min of entering each context was calculated. Ninety minutes after testing, rats were euthanized by decapitation to allow for maximal c-Fos expression (50). The left hemisphere of each brain was postfixed in 4% paraformaldehyde and used for immunohistochemistry and the right hemisphere was flash frozen and reserved for other experiments. Adrenals were weighed (see supplement).

### Immunohistochemistry & Microscopy

Sections including regions of interest were stained for c-Fos and sections including the dorsal and ventral hippocampus were double stained for doublecortin (DCX) and c-Fos. Region of interest included the medial frontal cortex (anterior cingulate cortex: ACC, prelimbic cortex: PRL, infralimbic cortex: IL), the nucleus accumbens (nucleus accumbens core: NAC, nucleus accumbens shell: NAS), the amygdala (basolateral amygdala: BLA, lateral amygdala: LA, central amygdala: CeA), the dorsal hippocampus (dHPC; dentate gyrus: dDG, cornu amonis 3: dCA3, cornu amonis 2: dCA2, cornu amonis 1: dCA1), and the ventral hippocampus (vHPC; vDG, vCA3, vCA1) (Fig. 2). These regions were chosen because they are implicated in negative cognitive bias or exhibit stress-related dysfunction (21,22,27,51). See supplemental methods for stereotaxic coordinates of each region and detailed staining protocols.

**Figure 1.**
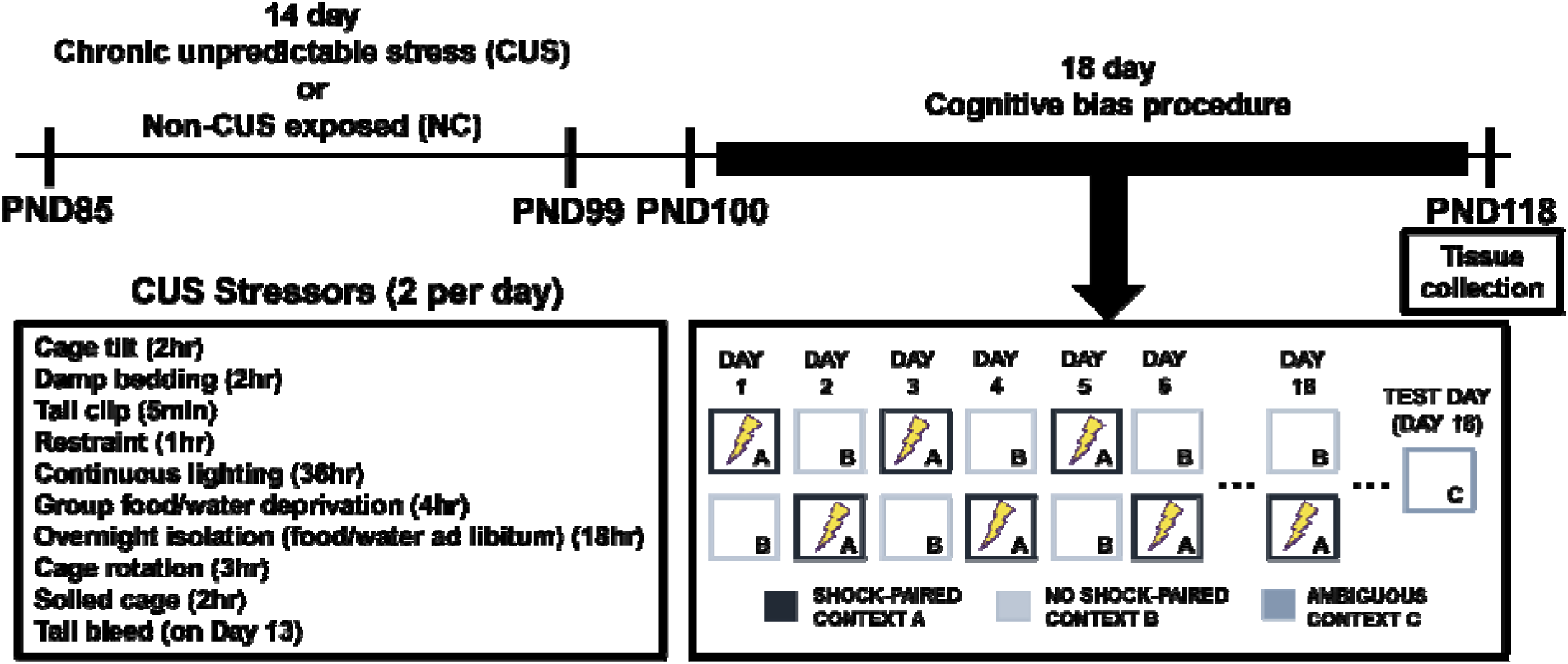
Experimental timeline. Rats were weighed on PND85, PND100 and PND118. PND: postnatal day.

**Figure 2.**
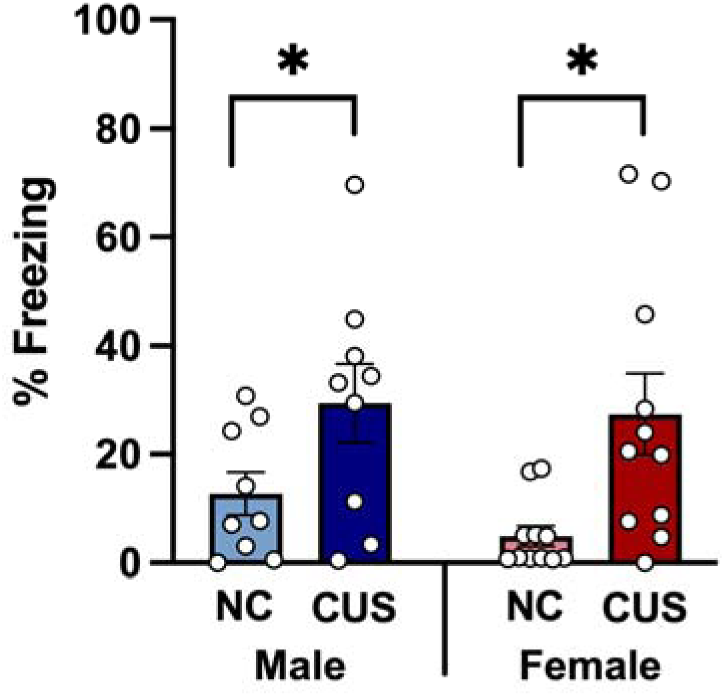
Percent freezing in the ambiguous context on test day. Mean (±SEM), * indicates p<0.05. NC: non-CUS-exposed, CUS: chronic unpredictable stress-exposed.

Images of each section stained for c-Fos were taken with a 40x objective lens on an Axio Scan.Z1 slide scanner (Carl Zeiss Microscopy, Thornwood, NY, USA). Areas of each region were calculated using FIJI/ImageJ software (52). For each region, c-Fos immunoreactive (c-Fos+) cells were summed across 4 sections per animal and then divided by the total area to calculate c-Fos cell density.

DCX-expressing cells were counted in 4 sections (2 dorsal, 2 ventral) per animal under 40x magnification (Nikon Eclipse 80i). For each animal, the density of DCX-expressing cells in each subregion was calculated by dividing total DCX-expressing cells by total area.

Images of each section stained for DCX and c-Fos were obtained using the Axio Scan.Z1 slide scanner (Carl Zeiss Microscopy, Thornwood, NY, USA). Two hundred and sixty DCX-expressing cells per animal were assessed for co-expression of c-Fos.

### Statistical analyses

We used two-way analysis of variance (ANOVA) with sex (male, female) and stress (NC, CUS) as between-subject factors to analyze freezing behavior in the ambiguous context. Repeated measures ANOVAs were used to assess c-Fos expression between subregions (frontal cortex: ACC, PRL, IL; nucleus accumbens: NAC, NAS; amygdala: BLA, LA, CeA; dentate gyrus: dDG, vDG; CA3: dCA3, vCA3; CA1: dCA1, vCA1), and DCX or DCX/c-Fos expression between subregions (dDG, vDG). *Post-hoc* tests used Newman-Keuls comparisons. Any *a priori* comparisons were subjected to Bonferroni correction. Significance level of p<0.05 was used. These statistical analyses were performed using Statistica software (v. 9, StatSoft, Inc., Tulsa, OK, USA). Behavior prediction, linear modeling, and network analysis were performed using R (version 4.4.2) and are described in the Supplement.

### Network analysis & behavior prediction modeling

For each group, functional networks were constructed using a consensus approach across three correlation methods (Pearson, Kendall, and shrinkage) and 3 network construction methods (percolation, weighted, and persistence). To explore the organizational principles underlying each network, we applied graph theory to extract several measures of centrality and clustering from each network. For each network metric, nodes were ranked and ranks were averaged together to achieve consensus across methods. For this analysis we focused on eigenvector centrality (EV), which describes the influence of a node within the network by considering the connectivity of that node as well as the connectivity of each neighboring node. EV was calculated for each node in each network, with 1 being the most influential and 0 being the least influential. Network analysis was conducted using the ConsensusConnectR app (https://github.com/AJMcGovernLab/ConsensusConnectRPrototype, see supplement for details).

To examine the relationship between regional activation and freezing, linear modeling was performed as previously described in Lee et al., 2026 (53). See supplement for details.

## RESULTS

### CUS increased freezing in the ambiguous context of the cognitive bias task regardless of sex

CUS rats displayed greater freezing behavior in the ambiguous context compared to NC rats, regardless of sex (Fig. 2, main effect of stress: F(1,36)=11.822, p<0.002, p^2^=0.247). There were no other significant main or interaction effects (p>0.395).

### CUS reduced DCX expression in the ventral hippocampus compared to NC

CUS reduced the density of DCX-expressing cells in the ventral, but not dorsal DG, compared to NC rats regardless of sex (Fig. 3, ventral DG p=0.001; dorsal DG p=0.730; region by stress interaction: F(1,35)=10.739, p=0.002, ^2^=0.235). There were no other significant main or interaction effects (p>0.180).

**Figure 3.**
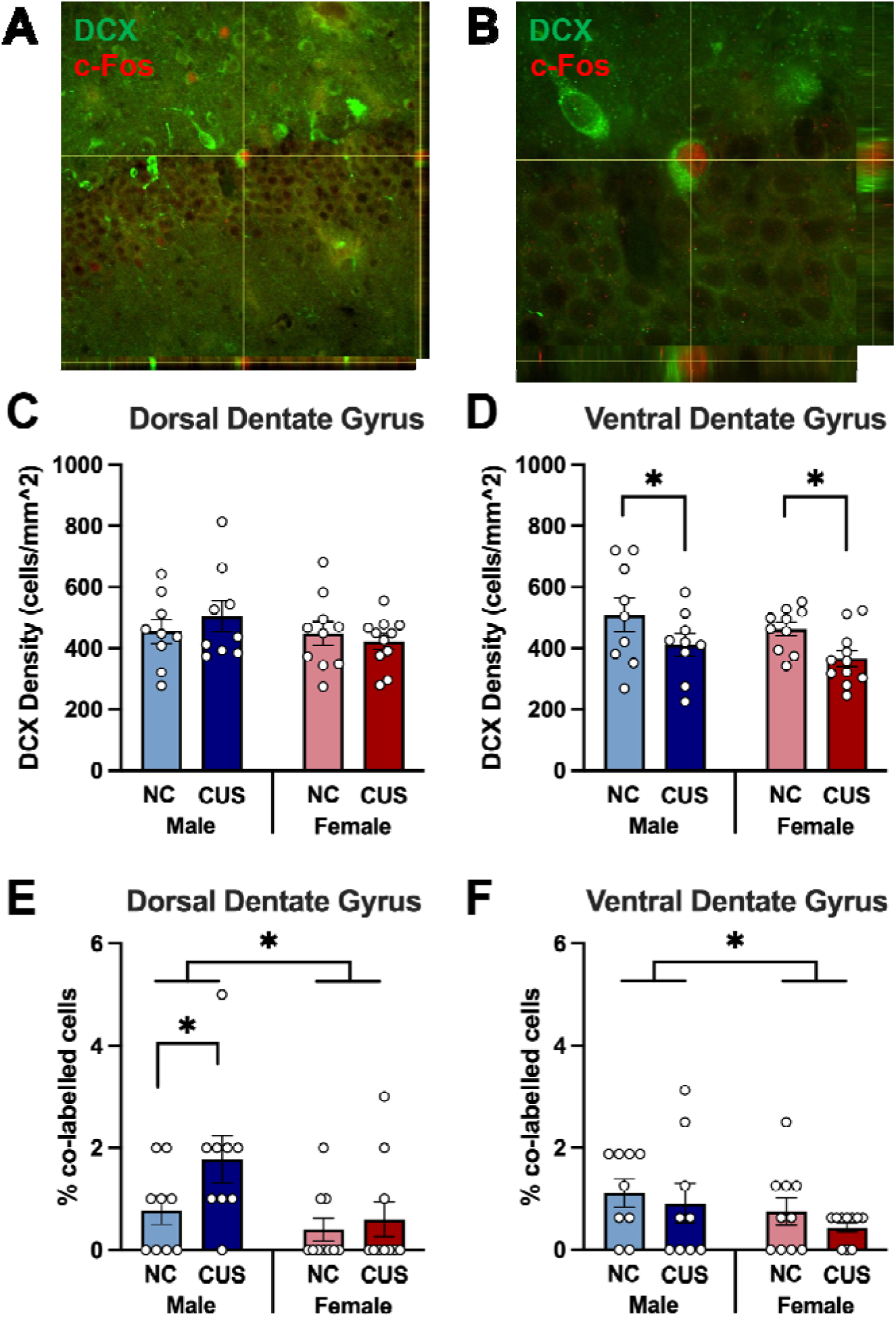
DCX/c-Fos immunohistochemical staining, DCX density, and % DCX/c-Fos colabeling. Representative images of DCX/c-Fos stain at 20x (A) and 60x (B) show co-localization of DCX and c-Fos in x-, y-, and z-planes. Density of DCX-expressing cells in the dorsal (C) and ventral (D) dentate gyrus and the percentage of DCX-expressing cells co-labelled with c-Fos in the dorsal (E) and ventral (F) dentate gyrus. Bar graphs show mean (±SEM). * indicates p<0.05. DCX: doublecortin, NC: non-CUS-exposed, CUS: chronic unpredictable stress-exposed.

### Males had greater activation of immature neurons in response to the ambiguous context than females

CUS males had a higher percentage of activated DCX-expressing cells (DCX co-labeled with c-Fos) in the dorsal DG compared to all other groups (all p’s<0.024; region by sex by stress interaction: F(1,34)=6.829, p=0.013, _p_^2^=0.167). There was also a significant main effect of sex (F(1,34)=7.117, p=0.012, ^2^=0.173). There were no other significant main or interaction effect (p>0.069).

### All rats had higher c-Fos expression in the IL compared to the PRL, and higher c-Fos expression in the PRL compared to the ACC

All rats, regardless of sex and stress condition, had higher c-Fos expression in the IL compared to the PRL (Fig. 4 D-F, p=0.001), and in the PRL compared to the ACC (p=0.005; main effect of region: F(2,58)=17.824, p=0.000001, ^2^=0.381). There were no other significant main or interaction effects (p>0.272).

**Figure 4.**
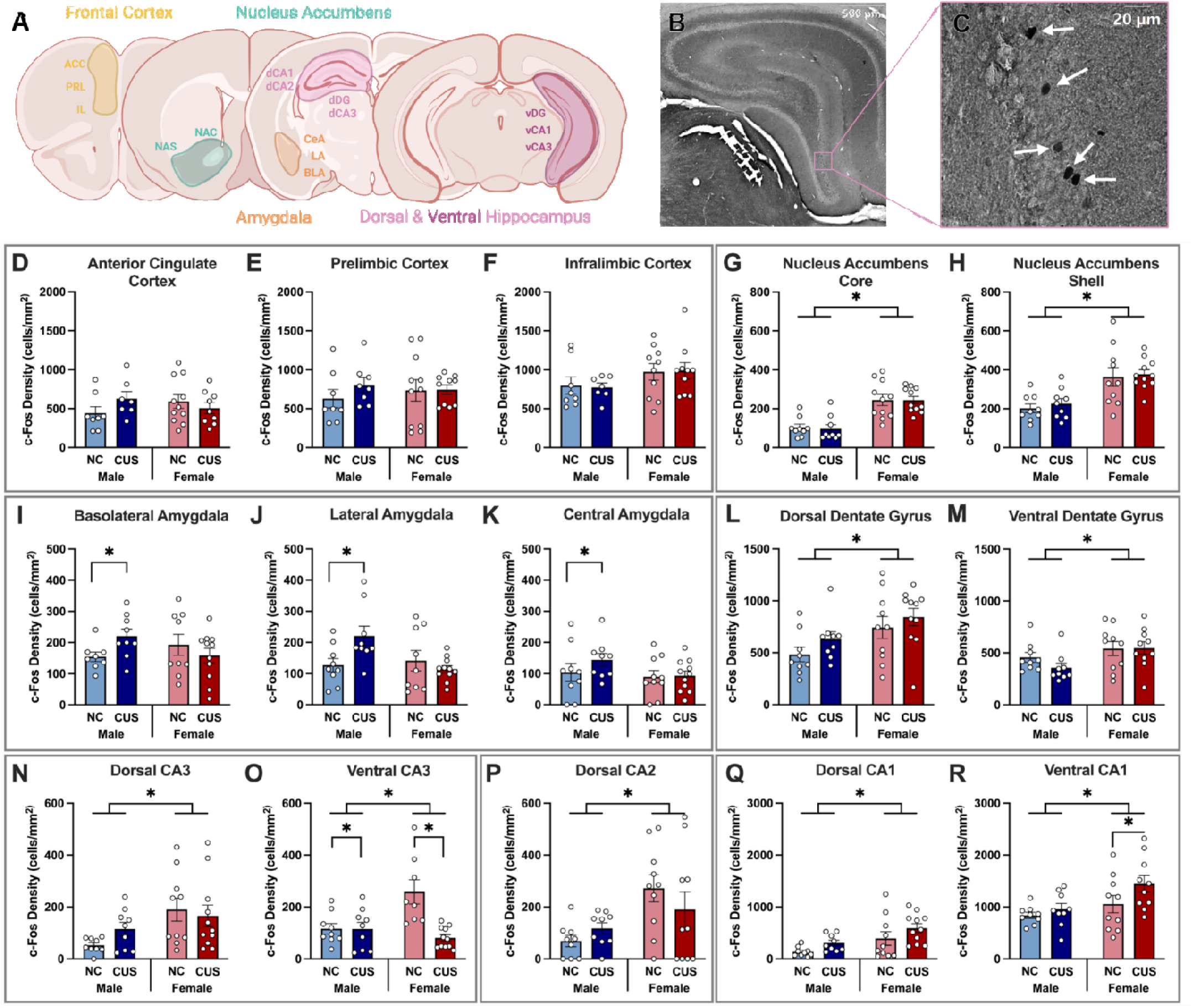
c-Fos immunohistochemical staining and c-Fos cell densities per region of interest. Regions of interest (A) including the frontal cortex (ACC: anterior cingulate cortex, PRL: prelimbic cortex, IL: infralimbic cortex), nucleus accumbens (NAC: nucleus accumbens core, NAS: nucleus accumbens shell), amygdala (BLA: basolateral amygdala, LA: lateral amygdala, CeA: central amygdala), dorsal hippocampus (dHPC; dDG: dorsal dentate gyrus, dorsal cornu ammonis 3: dCA3, dorsal cornu ammonis 2: dCA2, dorsal cornu ammonis 1: dCA1), and ventral hippocampus (vHPC; vDG: ventral dentate gyrus, vCA3: ventral cornu ammonis 3, vCA1: ventral cornu ammonis 1) were stained for c-Fos and imaged at 40x (B-C). White arrows in pop-out (C) indicate c-Fos+ cells. c-Fos cell densities were averaged across 4 sections containing each subregion for each animal. Bar graphs show mean (±SEM) density of c-Fos+ cells in subregions of the frontal cortex (D-F), nucleus accumbens (G-H), amygdala (I-K), the dorsal and ventral parts of the dentate gyrus (L-M), CA3 (N-O), CA2 (P), and CA1 (Q-R). Bar graphs show mean (±SEM). * indicates p<0.05. NC: non-CUS-exposed, CUS: chronic unpredictable stress-exposed.

### Females had higher c-Fos expression in the nucleus accumbens compared to males

Females had higher c-Fos expression in the nucleus accumbens compared to males (Fig. 4 G-H, main effect of sex: F(1,34)=37.692, p=0.000001, ^2^=0.526), regardless of region and stress condition. All rats had higher c-Fos expression in the NAS compared to the NAC (main effect of region: F(1,34)=122.467, p<0.000001, p^2^=0.779). There were no other significant main or interaction effects (p>0.167).

### CUS increased c-Fos expression in the amygdala in males only

CUS males had higher c-Fos expression in the amygdala than all other groups (Fig. 4 I-K, p’s<0.039), regardless of region. CUS did not influence amygdala c-Fos expression in females (p=0.695; sex by stress interaction: F(1,33)=5.902, p=0.021, □p^2^=0.152). All rats, regardless of sex and stress condition, had higher c-Fos expression in the BLA than in the LA (p=0.038), and in the LA than in the CeA (p=0.006; main effect of region: F(2,66)=12.218, p=0.00003, □p^2^=0.27). There were no other significant main or interaction effects (p>0.087).

### Females had higher c-Fos expression in all regions of the hippocampus compared to males; CUS increased activation of the CA1 in females only

Females had higher c-Fos expression in all analysed regions of the hippocampus (DG, CA3, CA2, CA1) compared to males (Fig. 4 L-R, main effects of sex: DG: F(1,30)=19.7560, p=0.0001, ^2^=0.397; CA3: F(1,32)=21.707, p=0.00005, □^2^=0.404; CA2: F(1,34)=8.393, p=0.007, □^2^=0.198; CA1: F(1,34)=9.418, p=0.004, □^2^=0.217), regardless of region or stress condition.

CUS rats had higher c-Fos expression in the CA1 region compared to NC rats (main effect of stress group: F(1,34)=4.711, p=0.037, □^2^=0.122), however, *a priori* comparisons revealed that this effect was driven by increased c-Fos expression in the vCA1 in CUS females (p=0.0006, Cohen’s d=0.753; CUS males: p=0.281). In contrast, in the vCA3, CUS rats had lower c-Fos expression than NC rats, which was not found in the dCA3 (region by stress group interaction: F(1,32)=5.38, p=0.027, □^2^=0.144).

Regionally, all groups also had higher c-Fos expression in the dDG compared to the vDG (main effect of region: F(1,30)=26.3410, p=0.00002, □_p_^2^=0.468) and higher c-Fos expression in the vCA1 compared to the dCA1, (main effect of region: F(1,34)=181.497, p<0.000001, □^2^=0.842). There were no other significant main or interaction effects across regions (p>0.099).

### CUS reduced edges, density, and clustering regardless of sex, but changes to global efficiency, average path length, and modularity were sex-specific

Using correlated c-Fos densities between subregions (Supplemental Fig. 3), we performed a network analysis using a consensus approach across methods, which revealed unique functional connectivity networks in each group (Table 1, Fig. 5, Supplemental Table 3). Regardless of sex and stress, all networks exhibited small-world properties. However, CUS decreased edges, density, and clustering in both sexes, resulting in sex-specific functional network reorganization. In males, CUS increased average path length with little change in modularity, suggesting some preservation of overall network architecture but an expansion of the network resulting in a decrease in global efficiency. In contrast, CUS decreased average path length and increased modularity but did not change global efficiency in females, suggesting increased network segregation into distinct functional communities that maintain their ability to communicate efficiently.

**Figure 5.**
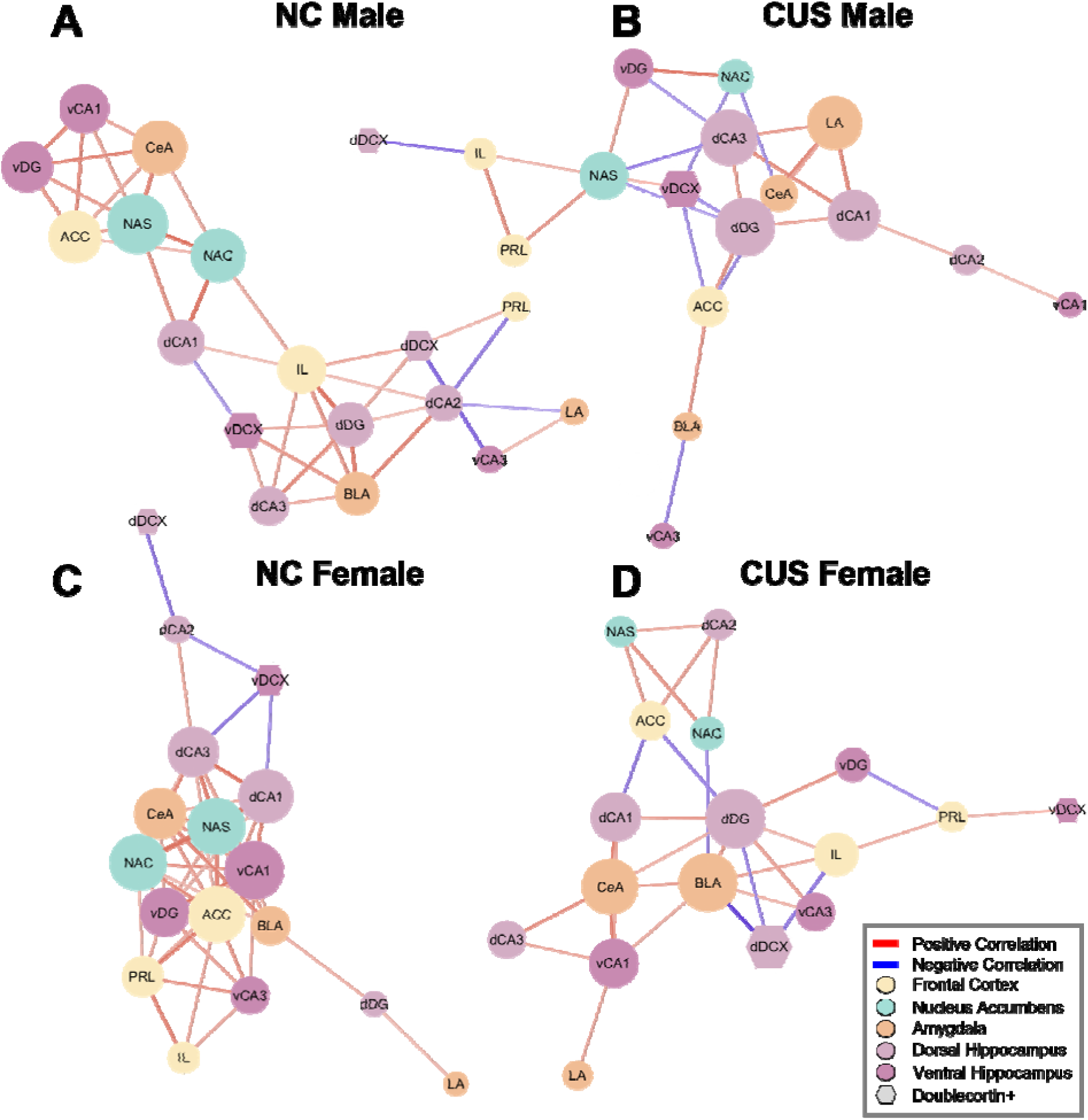
Visualization of force-directed functional connectivity networks constructed using Pearson correlations for NC males (A), CUS males (B), NC females (C), and CUS females (D). Positive correlations shown in red and negative correlations shown in blue, with brightness of color and weight of line corresponding to the strength of the correlation. Nodes are color coded based on brain area (yellow: frontal cortex, green: nucleus accumbens, orange: amygdala, light purple: dorsal hippocampus, dark purple: ventral hippocampus) and hexagonal nodes denote immature, doublecortin-expressing cells colabelled with c-Fos. NC: non-CUS-exposed, CUS: chronic unpredictable stress-exposed, ACC: anterior cingulate cortex, PRL: prelimbic cortex, IL: infralimbic cortex, NAC: nucleus accumbens core, NAS: nucleus accumbens shell, BLA: basolateral amygdala, LA: lateral amygdala, CeA: central amygdala, dDG: dorsal dentate gyrus, dCA3: dorsal cornu ammonis 3, dCA2: dorsal cornu ammonis 2, dCA1: dorsal cornu ammonis 1, dDCX: doublecortin-expressing cells in the dorsal dentate gyrus, vDG: ventral dentate gyrus, vCA3: ventral cornu ammonis 3, vCA1: ventral cornu ammonis 1, vDCX: doublecortin-expressing cells in the ventral dentate gyrus.

**Table 1.**
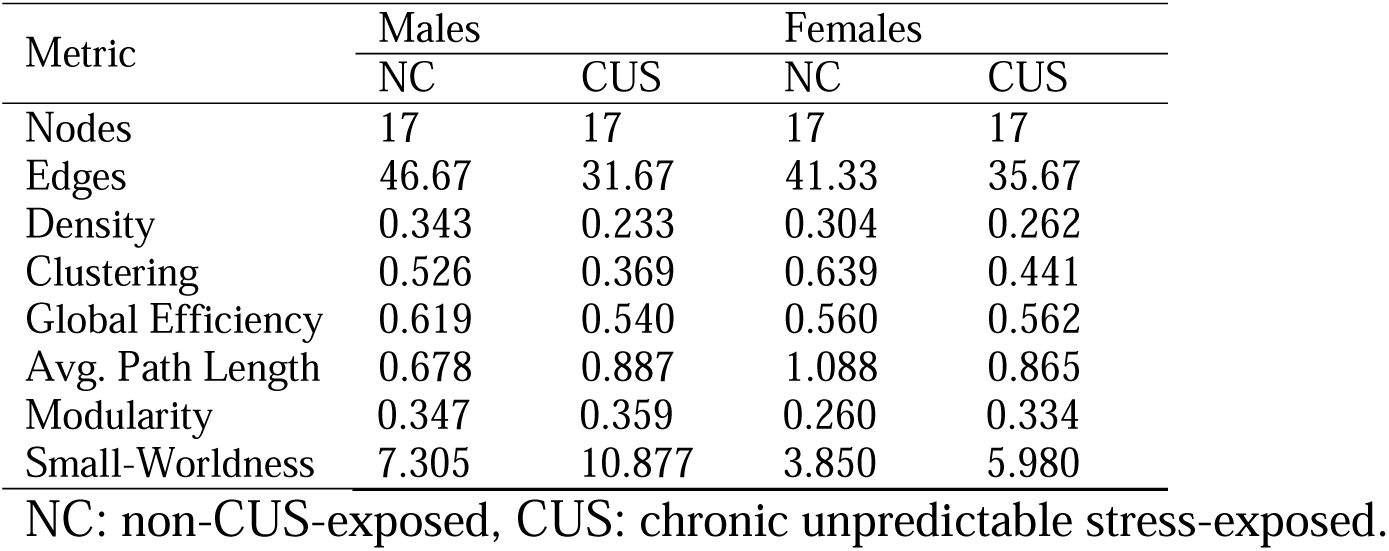
Global consensus network metrics from each group’s functional connectivity network.

### CUS reduced eigenvector centrality in subregions of the nucleus accumbens and the ventral hippocampus regardless of sex. In the amygdala, frontal cortex and in DCX-expressing cells, CUS had opposing effects on eigenvector centrality across sex

Our analysis revealed both stress-specific and sex-specific patterns in how CUS alter regional eigenvector (EV) centrality, which quantifies each region’s importance within th network based on both its connections and the importance of its neighbors (Fig. 6, Supplemental Table 3).

**Figure 6.**
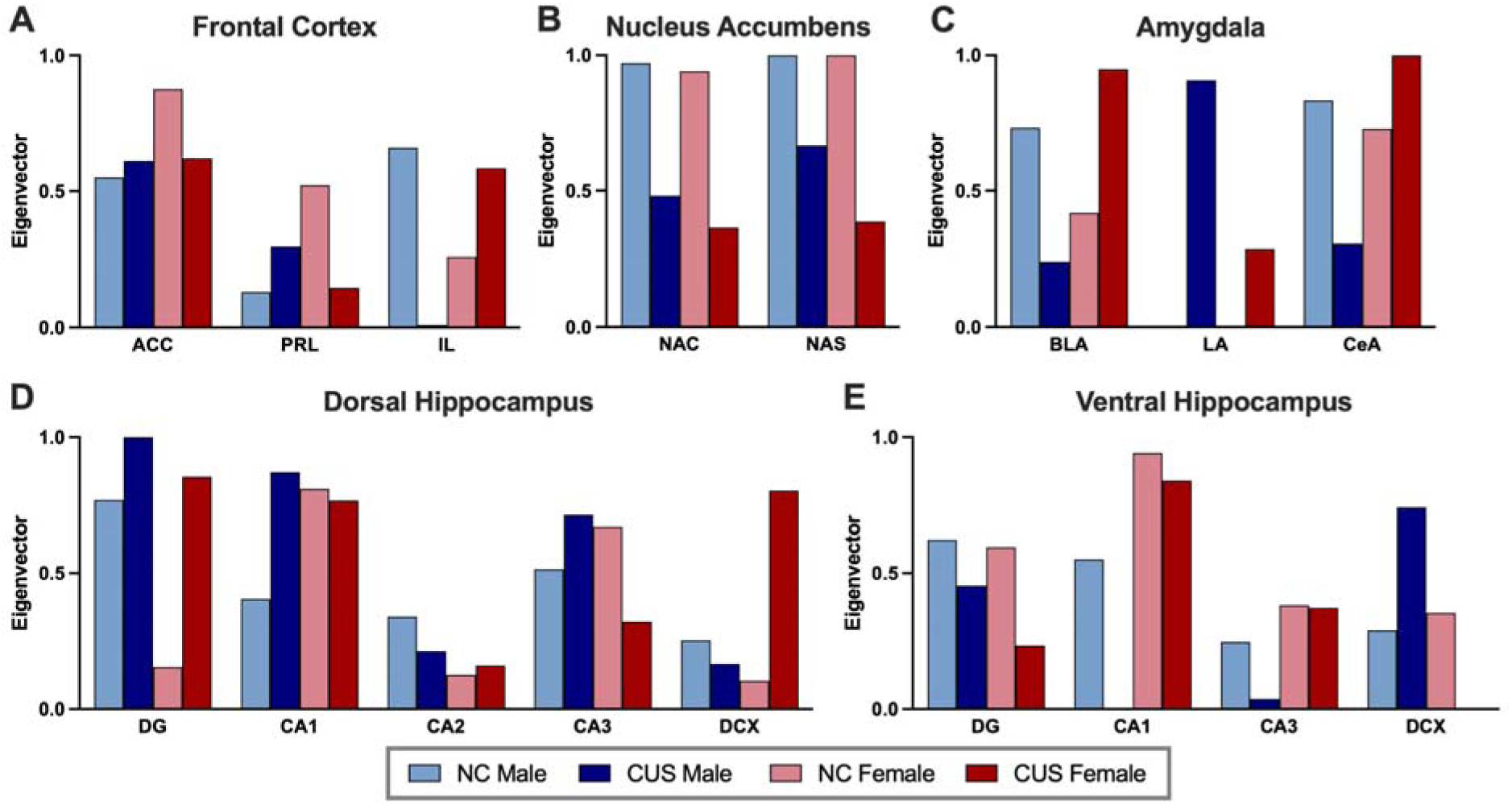
Eigenvector centrality (EV) of each subregion in the frontal cortex (A), nucleus accumbens (B), amygdala (C), dorsal hippocampus (D), and ventral hippocampus (E). EV ranges from 0 to 1 with 0 being the least “influential” in the network and 1 being the most. NC: non-chronic unpredictable stress-exposed, CUS: chronic unpredictable stress-exposed, ACC: anterior cingulate cortex, PRL: prelimbic cortex, IL: infralimbic cortex, NAC: nucleus accumbens core, NAS: nucleus accumbens shell, BLA: basolateral amygdala, LA: lateral amygdala, CeA: central amygdala, DG: dentate gyrus, CA3: cornu ammonis, CA2: cornu ammonis 2, CA1: cornu ammonis 1, DCX: doublecortin-expressing cells.

CUS had opposing effects across sex in most regions, with increased EV in females and decreased EV in males in the IL, amygdala (BLA, CeA), and dorsal hippocampus (CA2, DCX). CUS increased EV in males but decreased EV in females in the frontal cortex (ACC, PRL) and the hippocampus (dCA1, dCA3, vDCX). There were also substantial sex differences in EV noted across NC groups in all frontal regions, the BLA and most hippocampal regions.

CUS decreased EV, regardless of sex, in both the shell and core of the nucleus accumbens and in the DG and CA1 of the ventral hippocampus. However, CUS increased EV in the dorsal DG and LA in both sexes.

These differential patterns of regional influence suggest sex-specific neurobiological mechanisms underlying stress responses, with distinct brain regions making varying contributions to network activation and likely functional outcomes.

### Activity in the vCA1 paired with the amygdala or frontal regions drives the majority of group differences in freezing prediction

To examine the differences between groups in how an individual region or region pairs correlate with freezing, we produced linear models to test for interactions between sex and stress. No significant group interactions were identified for individual regions predictive of freezing. However, four region pairs were identified as having significant group differences, three of which include the vCA1: vCA1-LA, vCA1-ACC, vCA1-dDCX, and CeA-ACC (all p’s<0.037, Figure 7, Supplemental Table 7).

**Figure 7.**
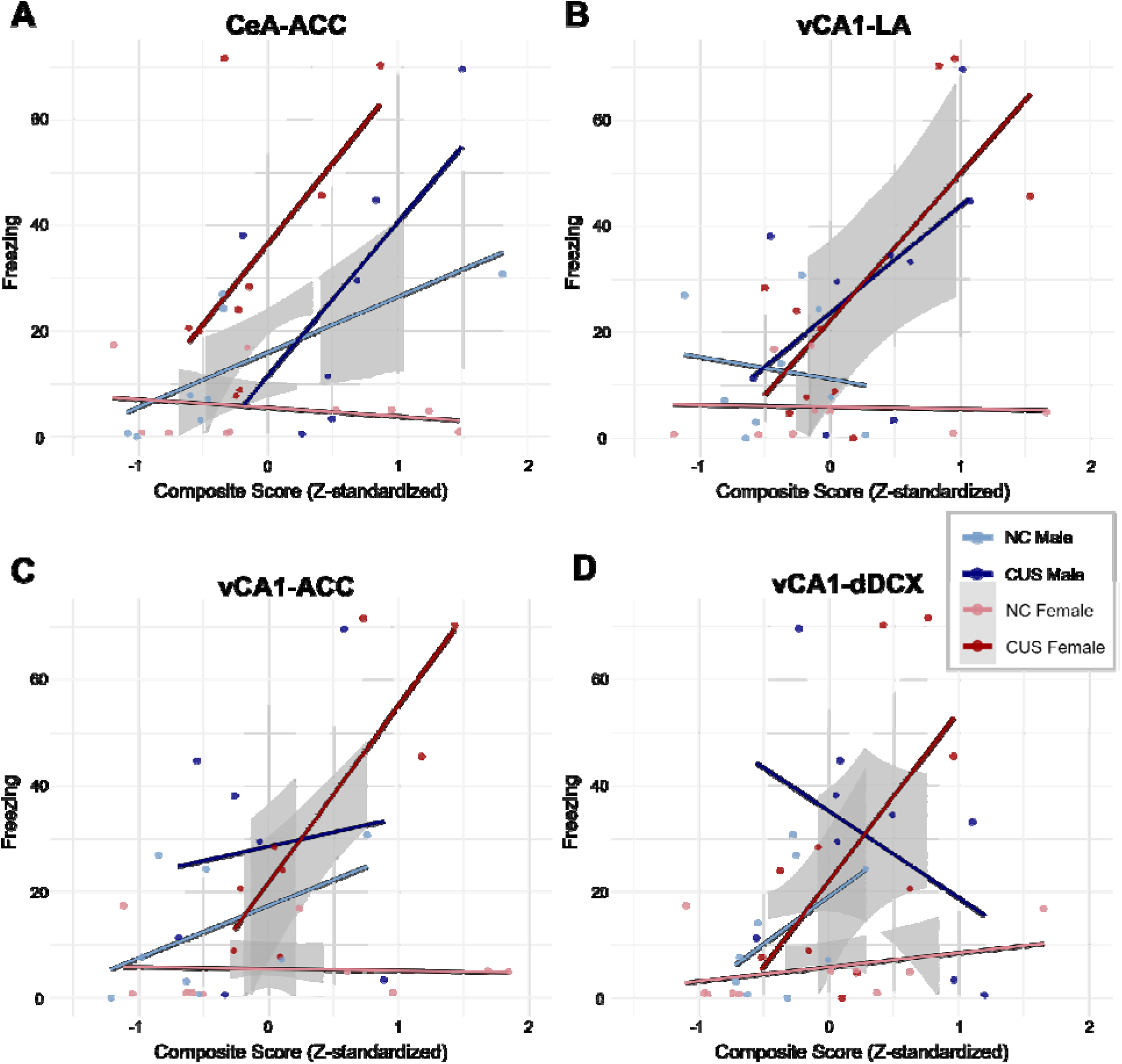
Correlations between freezing and four region pairs that exhibited significant group interactions in the linear model: CeA-ACC (A), vCA1-LA (B), vCA1-ACC (C), vCA1-dDCX (D). Grey shading indicates the 95% confidence interval of the group-specific linear fit. ACC: anterior cingulate cortex, LA: lateral amygdala, CeA: central amygdala, dDCX: DCX-expressing cells in the dorsal dentate gyrus, vCA1: ventral cornu ammonis 1, NC: non-CUS-exposed, CUS: chronic unpredictable stress-exposed.

Associations between freezing and activation were uniformly strengthened by CUS in females across these region pairs, whereas different patterns were seen in males exposed to CUS. In contrast to the strengthening of the positive correlation between vCA1-dDCX and freezing seen in CUS females compared to NC females, in males, CUS reversed the relationship, such that there was a negative correlation in CUS males. Intriguingly, this opposing relationship suggests that the activation of new neurons may support pattern separation differently across the sexes after stress exposure.

## DISCUSSION

Regardless of sex, CUS increased negative cognitive bias and decreased neurogenesis in the ventral dentate gyrus. Despite this, we found sex differences in neural activity signatures in response to the ambiguous context. We identified sex-specific patterns of neural activity including in new neurons, functional network efficiency, and regional contributions to the network, as well as the regional activity most predictive of freezing to the ambiguous context. Specifically, CUS increased activation of new neurons in the dorsal DG in males but not in females, yet CUS increased the importance of these new neurons in the network in females but not males. CUS increased amygdala activity in males but increased ventral CA1 activity in females in response to the ambiguous context, resulting in sex-specific functional network recruitment during the task. Regardless of stress exposure, females had more activity in the nucleus accumbens core and shell than males, yet CUS decreased the importance of these regions in the network in both sexes. Graph-based network analysis showed sex-specific functional reorganization resulting in a decrease in global efficiency in males and an increase in modularity but no change in global efficiency in females. Using linear modeling, we found that unique signatures of activity in pairs of regions including the ACC and ventral CA1 could predict freezing to the ambiguous context differentially by sex and stress exposure. Coordinated activity of the activated new neurons and the ventral CA1 positively predicted increased negative cognitive bias in females, with the reverse pattern seen in males. Taken together, these results suggest that CUS reorganizes the functional circuitry recruited during the cognitive bias task in a sex-specific manner, contributing to a growing body of evidence that sex-specific pathways may drive cognitive symptoms in MDD.

### CUS increased negative cognitive bias in both sexes

CUS increased freezing in the ambiguous context compared to controls in both sexes. Chronic stress exposure is a valuable tool to investigate depression neurobiology by reliably inducing depressive-like endophenotypes in rodents (54,55), and our results now include negative cognitive bias. Our findings are consistent with others using chronic stress paradigms and reward-based cognitive bias tasks in male rats (32,33,45–47), and female rats (32). However, these previously used tasks involve reward-seeking and require a motivated response from the animals to complete the task (56,57). As stress affects the salience of rewards/punishments and motivation (58), reward-based task design makes it difficult to interpret findings as simply negative cognitive bias. Our contextual fear-based cognitive bias task provides a robust, CUS-sensitive model of an important and understudied cognitive symptom of depression, without the need to disentangle anhedonia and avolition from negative cognitive bias.

### CUS reduced neurogenesis in the ventral DG regardless of sex but increased activation of DCX-expressing neurons in the dorsal DG in response to the ambiguous context only in males

Consistent with previous studies, CUS exposure reduced hippocampal neurogenesis in the ventral hippocampus of male and female rodents (59,60). Conversely, CUS increased activity of DCX-expressing neurons in males but not in females. Our findings are partially in line with work showing that immature neurons are activated during contextual fear retrieval in male and female rats (39,61). However, sex differences in disrupted activation patterns following CUS have not been previously explored. New neurons mature faster in male rats than in female rats (62), so additional research is required to determine if this sex difference is due to a shifted activity window in new neurons in females or other sex-specific mechanisms of integrating new neurons to be used during the cognitive bias task.

Our functional connectivity results also show that active new neurons contributed differently to male versus female networks after CUS exposure. CUS increased the importance of dorsal DCX-expressing cells in female networks and ventral DCX-expressing cells in male networks, but decreased the importance of ventral DCX-expressing cells in female networks, suggesting region- and sex-specific effects of CUS on functional integration of new neurons used to interpret ambiguity. Consistent with this result, our linear predictive modeling showed that coordinated activity in the ventral CA1-dorsal DCX region pair positively predicted freezing to the ambiguous context in females but negatively predicted freezing in males. This is consistent in part with previous work finding opposing patterns of correlations between ventral CA1 and dorsal DCX activity across sex in non-CUS-exposed rats that underwent a pattern separation task (39). Taken together, these results demonstrate sex-specific engagement of new neurons during pattern separation after stress exposure, warranting further investigation of sex-specific pattern separation mechanisms.

### CUS reorganized functional networks in a sex-specific manner, decreasing functional integration in males and increasing functional segregation in females

All networks exhibited small-world properties, regardless of sex and stress, suggesting the overall balance of functional integration and segregation is maintained following CUS. However, following CUS, we saw decreases in edges, density, and clustering in both male and female networks, suggesting some network fragmentation in both sexes. Further changes to average path length, modularity, and global efficiency were sex-specific. This functional network reorganization resulted in decreased global network efficiency in males, with no change to efficiency in females. In the human literature, comparisons of graph-based network properties between individuals with MDD and controls have yielded mixed results. One of the most consistent findings is the maintenance of small-world network properties across individuals with and without MDD (63–68), which parallels our findings in rats. The decrease in global efficiency seen in males following CUS is consistent with some resting-state fMRI studies of mixed-sex adult samples comparing individuals with MDD to controls (63,66,69). Two studies of mixed-sex adult samples that were predominantly female found an increase in global efficiency in individuals with MDD compared to controls (65,67), suggesting that network reorganization in MDD may be sex-specific. Given that we found sex-specific changes to average path length, modularity, and global efficiency, the heterogeneity in the literature may be due to insufficient accounting for sex differences in network changes.

The decrease in global efficiency seen in males following CUS indicates a decrease in functional integration that is not seen in females (70). Rather, female networks demonstrated an increase in modularity following CUS indicative of functional segregation (70), but as global efficiency did not decrease, communication between modules remained efficient. Although we found no sex differences in behavior, females have previously been reported to be more resilient to chronic stress-induced cognitive deficits than males, with stress exposure facilitating spatial performance in females but impairing it in males, despite evidence of dendritic atrophy in the CA3 region of both sexes. Similar findings are seen in postpartum females, with reduced hippocampal dendritic branching and neurogenesis associated with improved performance in a spatial radial arm maze (71,72). These, perhaps paradoxical, findings have been explained by a reduction in signal to noise ratio in females following stress exposure (72), which may correspond with an increase in functional segregation of the network without a loss of global efficiency as an adaptive stress response.

### The nucleus accumbens played a similar role in non-CUS-exposed male and female networks despite sex-specific activation patterns

Regardless of stress, activation of the nucleus accumbens core and shell was higher in females compared to males. However, the NAC and NAS seemed to fulfill similar roles between male and female networks. The NAC and NAS had very high EVs for both NC males and females, with CUS decreasing the EV of both regions in both males and females. The regional contribution analysis also identified the NAC and NAS as the only regions that contribute significantly to network similarity between non-CUS-exposed males and females (Supplemental Fig. 4B). Thus, although sex-specific activation patterns were seen across the nucleus accumbens, the subregions had similar roles in the NC networks, and CUS altered those roles in a sex-dependent manner.

### Activity in subregions of the amygdala and hippocampus predicted freezing in CUS males, whereas activity in the ventral CA1 predicted freezing in CUS females

Evidence from cFos densities, network analysis, and linear predictive modeling all converge to implicate CUS-induced network dysfunction involving the amygdala in males and the ventral CA1 in females in negative cognitive bias. First, we found that CUS increased amygdala activation in males and ventral CA1 activation in females in response to the ambiguous context. Our findings are partially consistent with other stress paradigms showing an increase in c-Fos expression in the amygdala of male mice (73) and in the ventral CA1 in both sexes (74). In the human literature, amygdala hyperactivation has been consistently associated with negative thinking and negative cognitive bias in people with MDD and other psychiatric disorders (22,75–78), although no studies have analyzed by sex. Furthermore, hippocampal activation is involved in context discrimination and contextual fear retrieval in a sex-specific manner (79–81). Thus, the increased negative cognitive bias we see in both sexes may be driven by chronic stress-induced dysfunction of amygdalohippocampal circuitry in both sexes that manifests in sex-specific c-Fos expression patterns.

In our linear models, CUS females exhibited positive correlations between freezing and activation of the ventral CA1 paired with activation of the LA, ACC, or dorsal DCX-expressing cells. In combined-sex or male-only samples, the ventral CA1 sends long-range projections to many stress-sensitive regions of the brain, including the medial prefrontal cortex and BLA, to communicate contextual fear memory information (82–86). In both the non-CUS-exposed male and female networks, we found that the ventral CA1 had a high EV and was densely connected within the network. Intriguingly, CUS had opposite effects on the role of the ventral CA1 in the male versus female networks, with the ventral CA1 becoming tightly connected to and exhibiting strong positive correlations with all three subregions of the amygdala, but not to subregions of the frontal cortex, in the CUS female network. This suggests that the CUS-induced increase in negative cognitive bias in females could be due to a loss of coordination between the frontal cortex, which is required for reappraisal of emotionally valanced information, and ventral CA1-BLA fear expression circuitry, which is consistent with the hypothesized circuit-level mechanism of negative cognitive bias in humans (21). These results indicate a need for further sex-specific characterization of the functional connectivity involved in negative cognitive bias, in both preclinical and clinical populations.

## CONCLUSIONS

Here we found that chronic unpredictable stress can be used to induce negative cognitive bias in both male and female rats. Our data show sex-specific CUS-induced changes in neurogenesis and functional connectivity involving multiple regions of the brain previously implicated in negative cognitive bias in humans and depressive-like endophenotypes in rodent models of MDD. CUS reorganized the functional networks recruited during cognitive bias testing, resulting in activity of the amygdala driving negative cognitive bias in males and activity of the ventral CA1 driving negative cognitive bias in females. A meta-analysis showed that sex is a potential moderator of differences in cognitive bias effect sizes (87), but to our knowledge, this is the first study to examine sex-specific neural mechanisms involved in negative cognitive bias in a rodent model of depression. These results underscore the importance of probing sex differences in depression neurobiology and investigating sex-specific targets for novel therapeutics, especially when treating cognitive symptoms.

## ACKNOWLEDGEMENTS

Financial support for this research was provided by an operating grant from the Natural Sciences and Engineering Research Council of Canada (2018–04301) to LAMG and a Canadian Institutes of Health Research project grant (PJT-186290) to LAMG, ABN, and TEH. ABN was supported by the Canadian Institutes of Health Research Masters and Doctoral Canada Graduate Scholarships. TEH was supported by the University of British Columbia Institute of Mental Health Marshalls Scholars Program. We thank Grace Y. Lee and Sophia H. Noh for helping with the behavioral testing of the animals.

