## Supplement Figures and Table captions for "Chronic unpredictable stress induces negative cognitive bias in rats with sex-specific changes in neurogenesis and functional connectivity"

**SUPPLEMENTAL METHODS**

**Supplemental Table 1**

Stressors included in the Chronic Unpredictable Stress (CUS) Paradigm

| Stressor | Duration | Description |
| --- | --- | --- |
| Cage tilt | 2 hr | Home cage tilted at 45°, water bottles removed. |
| Damp bedding | 2 hr | 200 ml tap water added to home cage. Rats moved to a dry and clean cage afterward. |
| Tail pinch | 5 min | Plastic clothes pin placed at the base of tail. |
| Restraint | 1 hr | Rats restrained in a well-ventilated plexiglass tube. |
| Continuous lighting | 36 hr | Colony lights on for 36 hr. |
| Group food/water deprivation | 4 hr | Rats deprived of food and water. |
| Overnight isolation (food/water ad libitum) | 18 hr |  |
| Cage rotation | 3 hr | Rats change cage partners for 3 hr. Rats returned to their original partner in a new clean cage after. |
| Soiled cage | 2 hr | Cagemates placed into a cage soiled by other rats. Rats returned to a new clean cage after. |
| Tail bleed | On day 13 of CUS procedure | Blood sample taken via tail vein. |
| Two stressors were presented daily, in a pseudorandom order, for 14 days. | | |

***Tissue preparation***

The left hemisphere of each brain was placed into a 4% paraformaldehyde solution for 24 h, and subsequently placed into a 30% sucrose in 0.1M phosphate buffered saline (PBS; pH 7.4) for another 24 h and then until sliced. Coronal sections (30µm) were sliced on a microtome and collected from approximately Bregma +3.72mm to -6.96mm (88). Sections were stored in an antifreeze solution (30% ethylene glycol, 20% glycerol in 0.1M phosphate buffer (PB; pH 7.4)) at -20 °C until immunohistochemistry assays were conducted.

***c-Fos immunohistochemistry***

Coronal sections were successively washed 4x in PBS for 10 min per wash and incubated at room temperature in a 0.6% hydrogen peroxide (H_2_O_2_; H1009, Sigma-Alrich, St. Louis, MO, USA) in distilled water (dH_2_O) for 30 min. Sections were then washed another 3x in 0.1M PBS for 10 min per wash, and then incubated at 4°C in c-Fos primary antibody (1:4000 Anti-c-Fos rabbit pAb; 190289; Abcam, Toronto, ON) for 24 hours. The next day, sections were washed 5x in 0.1M PBS for 10 min per wash and incubated overnight at 4°C in secondary antibody (biotinylated goat anti-rabbit IgG; 1:500; Vector Laboratories, Inc, Burlingame, CA). The last day, after another series of 5 washes in 0.1M PBS for 10 min per wash, sections were incubated in an avidin-biotin horseradish peroxidase solution (PK-4000, Vector Laboratories, Inc, Burlingame, CA) for 4 h at room temperature. Sections were washed 3x in 0.1M PBS for 10 min per wash and horseradish peroxidase was visualized using 3,3’ diaminobenzidine (DAB) in a 3 M sodium acetate buffer containing 2.5% nickel sulfate and 0.05% H_2_O_2_ (SK-4100, Vector Laboratories, Inc, Burlingame, CA) for 3 min. Sections were washed another 3x in 0.1M PBS for 10 min per wash and then mounted on Superfrost Plus slides (Fisher Scientific, Inc., Hampton, NH), let dry, dehydrated using increasing concentrations of ethanol (50%, 70%, 95%, 100% for 2, 2, 2, and 10 mins respectively), and then cleared with xylene for 10 min and coverslipped using Permount mounting medium (Fisher Scientific, Inc., Hampton, NH).

c-Fos protein immunostained brain images were taken with a 40x objective lens on an Axio Scan.Z1 slide scanner with Zen 3.0 software (Carl Zeiss Microscopy, Thornwood, NY, USA). Digital images of regions of interest included the frontal cortex (anterior cingulate cortex = ACC, prelimbic cortex = PRL, infralimbic cortex = IL) within bregma 3.72 mm and 2.52 mm, the nucleus accumbens (nucleus accumbens core = NAC, nucleus accumbens shell = NAS) within bregma 1.92 mm and 0.96 mm, the amygdala (basolateral amygdala = BLA, lateral amygdala = LA, central amygdala = CeA) within -2.16 mm and -2.92 mm, the dorsal hippocampus (dHPC; dentate gyrus (dDG), dCA3, dCA2, dCA1) within -2.64 mm and -4.56 mm, and the ventral hippocampus (vHPC; vDG, vCA3, vCA1) within -5.76 mm and -6.36 mm were taken using 4x and 10x objectives. Photomicrographs were used to trace an outline of each subregion of interest to calculate the area of each region using ImageJ/FIJI software (2). Cell counts of c-Fos immunoreactive (ir) cells were conducted by experimenters’ blind to experimental condition and averaged across 4 sections per animal per subregion of interest. c-Fos-ir cell density for each subregion of interest was calculated by dividing the cell count by the corresponding area in mm^2^ for each animal.

***Doublecortin immunohistochemistry***

Coronal sections were stained for doublecortin (DCX) using previously established protocols (48). Tissue was washed 1x in distilled water and 3x in PBS for 10 min each before being incubated in 0.6% hydrogen peroxide in distilled water for 30 mins at room temperature. Sections were again washed 3x in PBS for 10 min each, then incubated in DCX primary antibody (1:1000 goat anti-doublecortin polyclonal IgG; SC-8066; Santa Cruz Biotechnology, Dallas, TX) for 24 hr at 4°C. Next, sections were washed 5x in PBS for 10 min each and then incubated in secondary antibody (1:500 biotinylated rabbit anti-goat IgG; BA-5000; Vector Laboratories, Burlingame, CA) overnight at 4°C. The next day, tissue was washed 5x in PBS for 10 min each. Sections were incubated in an avidin-biotin solution containing horseradish peroxidase (HRP) (PK-4000; Vector Laboratories, Burlingame, CA) for 4 hr at room temperature, then washed again with PBS 3x for 10 min each. Finally, to visualize DCX-expressing (DCX+) cells, sections were incubated for 3 min at room temperature in an HRP-reactive 3,3’-diaminobenzidine (DAB)/nickel substrate solution containing 2.5% nickel sulfate and 0.05% H2O2 (SK-4100; Vector Laboratories, Burlingame, CA). Tissue was washed another 3x in PBS for 10 min each then mounted on Superfrost Plus slides (Fisher Scientific, Hampton, NH). Slides were then dehydrated in ethanol prior to coverslipping.

DCX was visualized in the dorsal hippocampus (dHPC, bregma -2.92 to -4.56 mm) and ventral hippocampus (vHPC, bregma -5.76 to -6.36 mm) using a Nikon Eclipse 80i microscope. The locations of regions of interest were identified according to Paxinos and Watson, 2005 (1). DCX+ cells in the dentate gyrus (DG) in two sections of dHPC and two sections of vHPC per animal were manually counted using the 40x objective. Digital images of the sections used for cell counts were acquired using the 10x objective. The outline of each DG was traced, including the granule cell layer but not the subgranular zone, and area calculated using ImageJ/FIJI software (2). Finally, the density of DCX+ cells in each subregion was calculated by summing each animal’s cell counts and areas, then dividing total DCX+ cells by total area in mm^2^ for both the dDG and vDG.

***c-Fos/DCX double labelling***

To determine whether DCX+ cells also co-expressed the activity marker, cFos, a second series of hippocampal sections was stained for both DCX and cFos. The tissue was washed 4x in PBS for 7.5 min each then incubated in a solution containing 10% Triton-X in PBS for 30 min at room temperature. Sections were then incubated in blocking solution containing 10% normal donkey serum (NDS) and 0.3% Triton-X in PBS for 1 hr at room temperature. Next, the tissue was transferred to primary antibody solution with antibodies for DCX (1:1000 goat anti-doublecortin polyclonal IgG; SC-8066; Santa Cruz Biotechnology, Dallas, TX) and cFos (1:1000 rabbit anti-cFos polyclonal IgG; ab190289; Abcam, Cambridge, UK), diluted with the same blocking solution previously described. After incubating for 48 hr at 4°C, sections were washed in PBS 3x for 10 min each at room temperature, then transferred to secondary antibody solution prepared in the same blocking solution previously described containing Alexa Fluor 488 (1:500 donkey anti-goat IgG; A-11055; Invitrogen, Waltham, MA) and Alexa Fluor 594 (1:500 donkey anti-rabbit IgG; A-21207; Invitrogen, Waltham, MA). Tissue was incubated in secondary antibody solution for 4 hr at room temperature and all procedures from this point on occurred in the dark. Sections were again washed 3x in PBS for 10 min each at room temperature. Next, sections underwent a 2.5 min DAPI incubation (1:5000 in PBS; D1306; Life Technologies, Etobicoke, ON) at room temperature followed by 3x 5 min PBS washes. Tissue was kept at 4°C in PBS until being mounted on Superfrost Plus slides (Fisher Scientific, Hampton, NH). Once dry, slides were coverslipped with PVA-DABCO.

Digital fluorescent images of each section were obtained using the Axio Scan.Z1 slide scanner with Zen 3.0 software (Carl Zeiss Microscopy, Thornwood, NY, USA). Two hundred and sixty DCX+ cells were manually assessed for co-expression of c-Fos across the dorsal and ventral regions (2+ sections per region). Percentages of double-labelled cells for each animal were calculated separately by dorsal versus ventral regions.

***Statistical analyses***

Outliers that fell >2 standard deviations from the mean were removed from analyses (Supplemental Table 2). Missing data were imputed for the network analysis using predictive mean matching (R package “mice”; m = 5, 10 iterations, seed 123).

| **Supplemental Table 2.** | | | | |
| --- | --- | --- | --- | --- |
| Number of data points removed or missing from c-Fos expression in each group due to being an outlier (>2 standard deviations away from the mean) or damaged tissue. | | | | |
|  | Male NC | Male CUS | Female NC | Female CUS |
| ACC | 1 | 2 | 0 | 2 |
| PRL | 1 | 1 | 0 | 1 |
| IL | 1 | 2 | 0 | 1 |
| NAC | 0 | 0 | 0 | 0 |
| NAS | 0 | 0 | 1 | 0 |
| BLA | 1 | 0 | 2 | 0 |
| LA | 0 | 0 | 2 | 0 |
| CeA | 0 | 0 | 1 | 0 |
| dDG | 0 | 0 | 1 | 0 |
| dCA3 | 0 | 0 | 1 | 0 |
| dCA2 | 0 | 0 | 1 | 0 |
| dCA1 | 0 | 0 | 1 | 0 |
| vDG | 0 | 0 | 1 | 1 |
| vCA3 | 0 | 0 | 3 | 1 |
| vCA1 | 0 | 0 | 1 | 1 |
| dDCX | 0 | 0 | 1 | 0 |
| vDCX | 0 | 0 | 1 | 0 |

NC: non-CUS-exposed, CUS: chronic unpredictable stress-exposed, ACC: anterior cingulate cortex, PRL: prelimbic cortex, IL: infralimbic cortex, NAC: nucleus accumbens core, NAS: nucleus accumbens shell, BLA: basolateral amygdala, LA: lateral amygdala, CeA: central amygdala, dDG: dorsal dentate gyrus, dCA3: dorsal cornu ammonis 3, dCA2: dorsal cornu ammonis 2, dCA1: dorsal cornu ammonis 1, dDCX: doublecortin-expressing cells in the dorsal dentate gyrus, vDG: ventral dentate gyrus, vCA3: ventral cornu ammonis 3, vCA1: ventral cornu ammonis 1, vDCX: doublecortin-expressing cells in the ventral dentate gyrus.

***Network construction and analysis***

Functional connectivity networks were constructed for each group using ConsensusConnectR (latest version: <https://github.com/AJMcGovernLab/ConsensusConnectRPrototype>). Following imputation of missing values, three correlation estimators were computed per group: Pearson, Kendall, and a shrinkage estimator. Each correlation matrix was used to construct networks under three approaches, giving nine method–approach combinations per group. The weighted approach retains all edges and uses the correlation value as the edge weight. The percolation approach raises the correlation threshold for each group in 0.01 increments until every region remains connected to at least one other. The persistence approach uses binary edges and tests the network at thresholds from 0.10 to 0.90 in steps of 0.05 and recomputes node metrics at each threshold, taking the area under the threshold × metric curve as the final value. For each node metric, nodes were ranked within each of the nine combinations (rank 1 = highest), ranks were averaged across combinations, and the averaged ranks were rescaled linearly to the interval 0–1, such that the highest-ranked region in a group scores 1 and the lowest-ranked scores 0. Global and nodal consensus metrics were calculated. Functional connectivity networks were constructed using the igraph package, where nodes represent brain regions and edges represent correlations. Force-directed network visualizations were created with Cytoscape (v3.10.1) (3).

Across groups, consensus networks were compared using a Jaccard similarity index and a regional contribution analysis. Jaccard similarity is computed as the number of edges shared by two networks divided by the number of edges present in either. 1 indicates identical edge sets and 0 describes no shared edges. The value reported for each network comparison is the mean across the nine method–approach combinations.

The regional contribution analysis determines whether a given node contributes to network similarity or difference between two groups by removing that node from both networks and examining the change in Jaccard similarity (contribution score). A positive score indicates the two networks became more similar after the node was removed, identifying that region as a source of difference between the groups. A negative score means the networks became less similar once it was removed, thus contributing to similarity between the networks. Significance was tested by permutation with randomly redistributed node labels, held consistent across methods, with p = (b + 1)/(m + 1) where b is the number of permutations with an absolute change at least as large as observed and m is the number of permutations. The number of permutations was set to the larger of 1000 or a value scaled to the number of tests with a safety factor of 2, so that the attainable p-value floor remains below the threshold needed after FDR correction. Each region is tested twice (A vs B and B vs A) by construction.

***Linear modeling and behavior prediction***

To investigate the relationship between neural activity and behavior in each group, we examined regional connectivity by analyzing c-Fos density across 15 brain regions (Fig. 4) and DCX+ cells in the dDG and vDG (dDCX, vDCX). We systematically tested each individual brain region and pairs of regions as predictors of freezing. Data was imported into R with packages tidyverse, glmnet, and corrplot loaded for data management, analysis, and visualization. Using the lm function in R for multiple linear regression, we modeled freezing as a dependent variable influenced by c-Fos expression in up to four brain regions. Adjusted R^2^ was extracted after each individual correlation calculation, which accounts for the number of model predictors.

To examine group differences in the brain region combinations most predictive of freezing, we introduced interaction terms to our linear regression models, including the main effects of group, region, and their interactions. A 2-way ANOVA was performed to test for statistically significant interactions between regions and group correlations with behavior.

**SUPPLEMENTAL RESULTS**

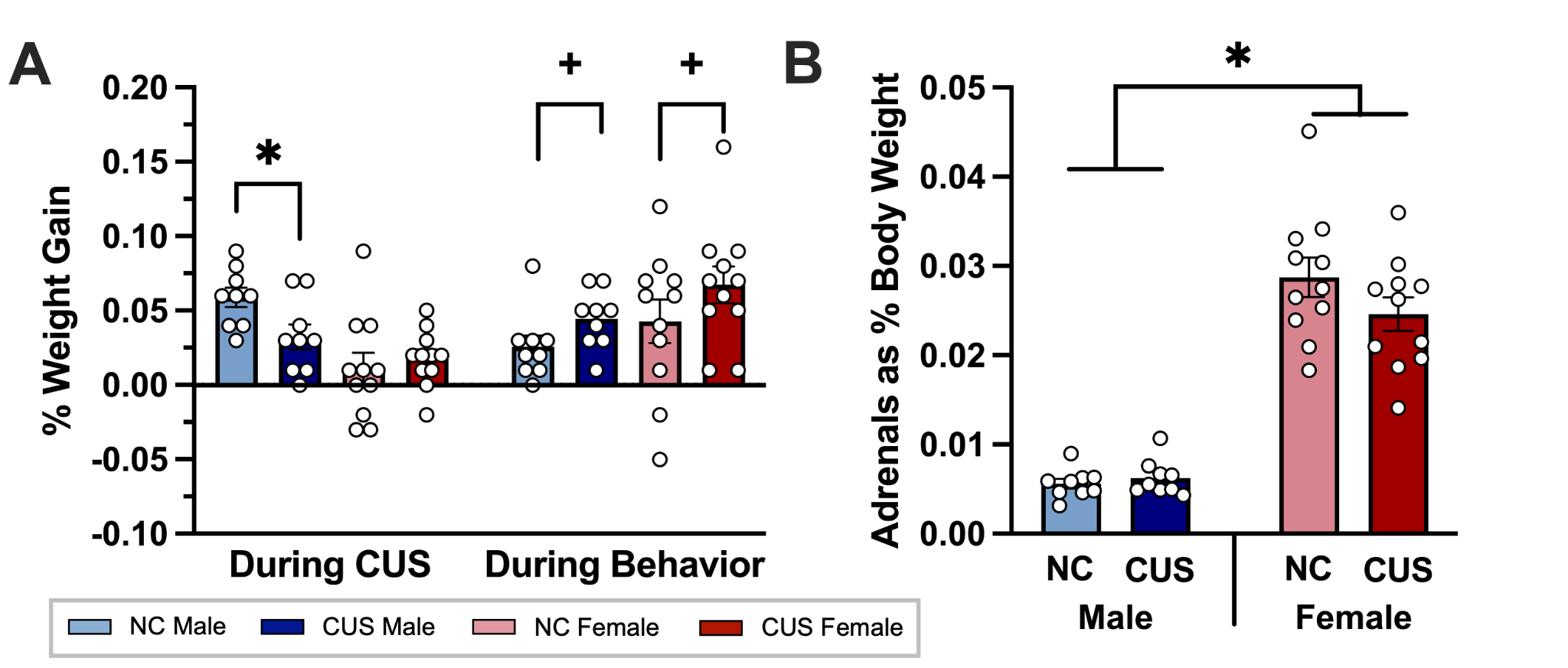

**Supplemental Figure 1. Percent weight gain during CUS (day 1 of CUS to day 1 of cognitive bias training) and behavior (day 1 of cognitive bias training to test day) (A) and adrenal mass as a percentage of body weight on test day (B).** NC: non-CUS-exposed, CUS: chronic unpredictable stress-exposed. Bar graphs show mean (+/- SEM). * indicates p<0.05, + indicates p=0.058.

***CUS males gained less weight than NC males after the 14-day CUS***

After CUS, CUS male rats had a lower percentage of weight gain than NC male rats (p=0.011), not found in females (p=0.116; sex by stress group interaction: F(1,33)=9.354, p=0.004, Ƞ_p_^2^=0.221). After the cognitive bias task, the percentage of weight gain did not significantly differ by sex or stress group (p’s>0.058). However, there was a near-significant effect of stress, such that CUS rats had a higher percentage of weight gain than NS rats, regardless of sex (main effect of stress: F(1,34)=3.852, p=0.058, Ƞp^2^=0.102).

When examining adrenal mass after the CUS and cognitive bias procedures, females had a higher adrenal mass as a percentage of body weight than males (main effect of sex: F(1,36)=157.825, p<0.001, Ƞp^2^=0.814), but there were no stress group or interaction effects (p’s>0.158).

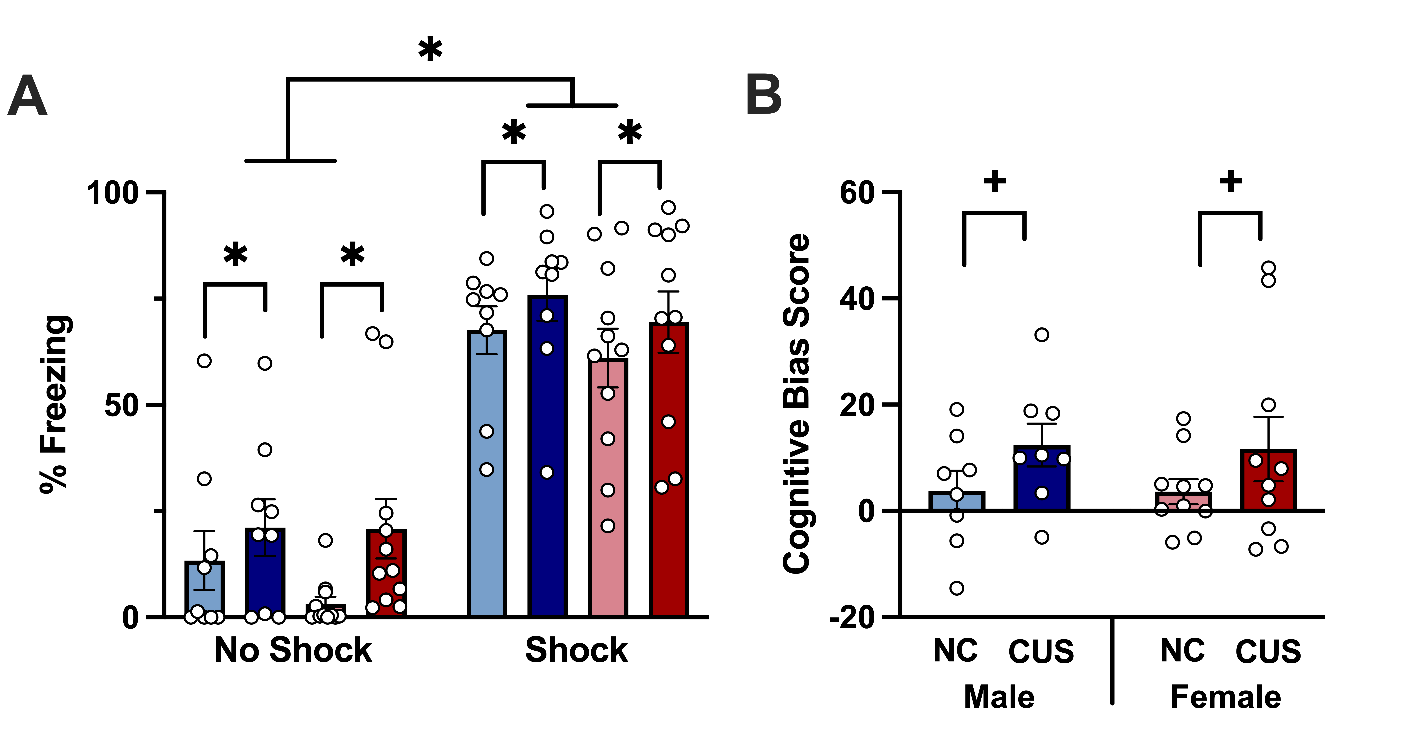

**Supplemental Figure 2. Percent time freezing in no shock and shock contexts on day 16 (A) and cognitive bias score on test day (B).** Cognitive bias score was calculated by subtracting freezing in the no shock context on day 16 from freezing in the ambiguous context on test day. A more positive score indicates a more negative cognitive bias. NC: non-CUS-exposed, CUS: chronic unpredictable stress-exposed. Bar graphs show mean (+/- SEM). * indicates p<0.05, + indicates p=0.069**.**

***CUS-exposed animals froze more across all contexts and had a greater fear response to the ambiguous context than non-CUS-exposed animals***

Regardless of sex and stress, all rats showed good discrimination of the contexts, freezing significantly more in the shock-paired context than the no shock-paired context on day 16 of training (main effect of context: F(1,36)=206.91, p<0.000001, Ƞp^2^=0.852, Supplemental Fig. 2). Moreover, CUS rats displayed more freezing than NS rats in both the shock-paired and no shock-paired contexts (Main effect of stress group: F(1,36)=4.316, p=0.045, Ƞp^2^=0.107), regardless of sex. All other effects p>0.257.

When examining the cognitive bias score, no effects were significant (p’s>0.069). However, there was a near-significant effect of stress group such that CUS rats had a higher negative cognitive bias than NS rats (near significant main effect of stress group: F(1,32)=3.553, p=0.069, Ƞp^2^=0.100), regardless of sex.

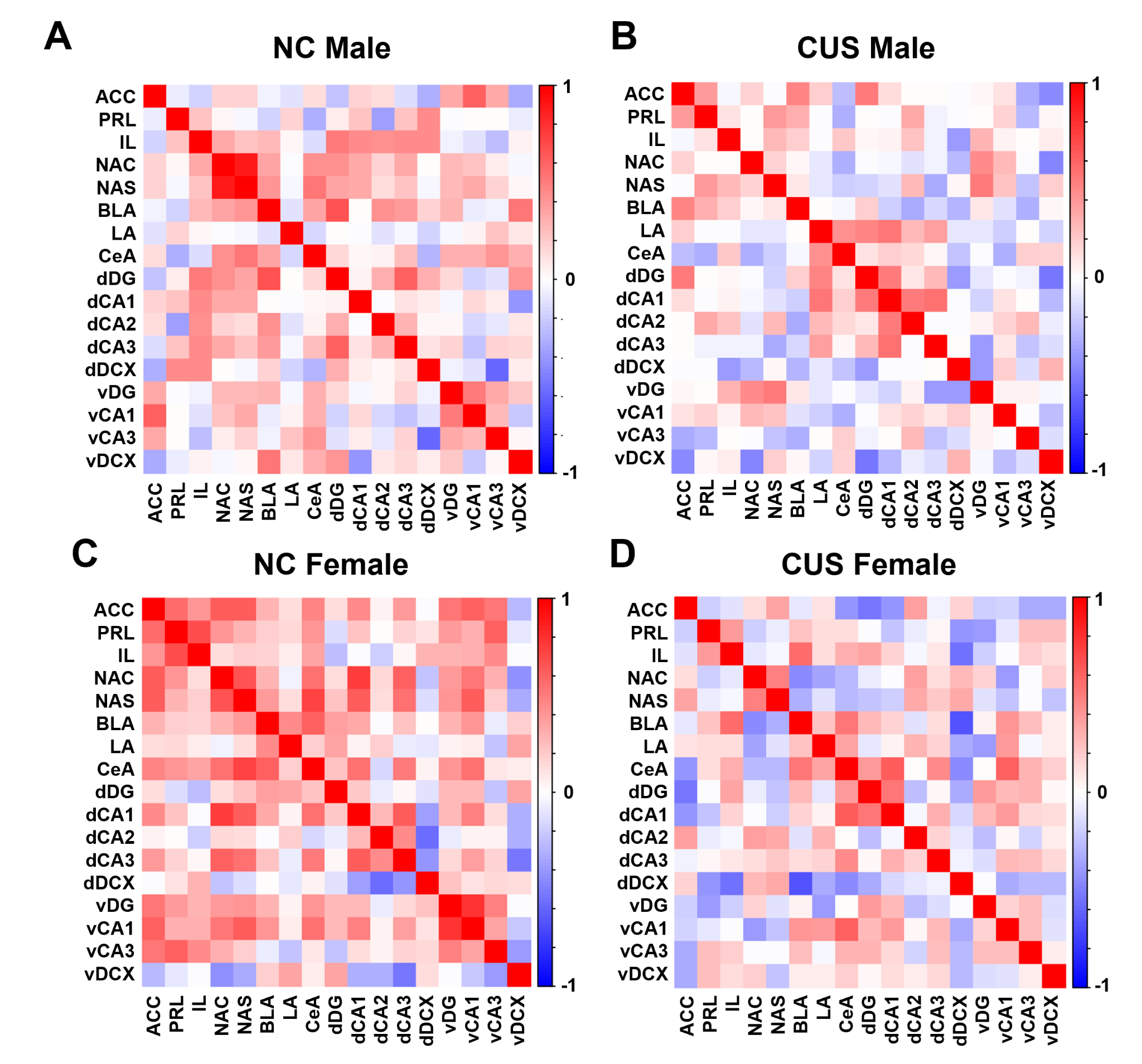

**Supplemental Figure 3. Consensus correlation matrices across each subregion included in the network analysis for NC males (A), CUS males (B), NC females (C), and CUS females (D).** Color gradient indicates the R-value for each correlation with a perfect positive correlation in red and a perfect negative correlation in blue. NC: non-CUS-exposed, CUS: chronic unpredictable stress-exposed, ACC: anterior cingulate cortex, PRL: prelimbic cortex, IL: infralimbic cortex, NAC: nucleus accumbens core, NAS: nucleus accumbens shell, BLA: basolateral amygdala, LA: lateral amygdala, CeA: central amygdala, dDG: dorsal dentate gyrus, dCA3: dorsal cornu ammonis 3, dCA2: dorsal cornu ammonis 2, dCA1: dorsal cornu ammonis 1, dDCX: doublecortin-expressing cells in the dorsal dentate gyrus, vDG: ventral dentate gyrus, vCA3: ventral cornu ammonis 3, vCA1: ventral cornu ammonis 1, vDCX: doublecortin-expressing cells in the ventral dentate gyrus.

**Supplemental Table 3**

Node centrality metrics from consensus networks.

| Region | Metric | Males | | Females | |
| --- | --- | --- | --- | --- | --- |
|  |  | NC | CUS | NC | CUS |
| ACC | Strength | 0.713 | 0.358 | 0.866 | 0.857 |
|  | Eigenvector | 0.676 | 0.469 | 0.813 | 0.748 |
| PRL | Strength | 0.103 | 0.193 | 0.480 | 0.086 |
|  | Eigenvector | 0.079 | 0.171 | 0.461 | 0.077 |
| IL | Strength | 0.699 | 0.651 | 0.236 | 0.393 |
|  | Eigenvector | 0.576 | 0.631 | 0.141 | 0.490 |
| NAC | Strength | 0.912 | 0.789 | 0.850 | 0.414 |
|  | Eigenvector | 0.899 | 0.730 | 0.922 | 0.308 |
| NAS | Strength | 1.000 | 0.679 | 1.000 | 0.500 |
|  | Eigenvector | 1.00 | 0.604 | 1.000 | 0.427 |
| BLA | Strength | 0.654 | 0.339 | 0.315 | 0.921 |
|  | Eigenvector | 0.532 | 0.306 | 0.375 | 0.909 |
| LA | Strength | 0.000 | 0.789 | 0.024 | 0.243 |
|  | Eigenvector | 0.000 | 0.784 | 0.008 | 0.238 |
| CeA | Strength | 0.846 | 0.156 | 0.575 | 1.000 |
|  | Eigenvector | 0.878 | 0.207 | 0.586 | 1.000 |
| dDG | Strength | 0.978 | 1.000 | 0.055 | 0.886 |
|  | Eigenvector | 0.727 | 1.000 | 0.125 | 0.790 |
| dCA3 | Strength | 0.404 | 0.596 | 0.630 | 0.243 |
|  | Eigenvector | 0.446 | 0.658 | 0.602 | 0.364 |
| dCA2 | Strength | 0.177 | 0.128 | 0.000 | 0.300 |
|  | Eigenvector | 0.158 | 0.126 | 0.000 | 0.224 |
| dCA1 | Strength | 0.213 | 0.725 | 0.811 | 0.686 |
|  | Eigenvector | 0.374 | 0.730 | 0.836 | 0.713 |
| vDG | Strength | 0.632 | 0.468 | 0.614 | 0.357 |
|  | Eigenvector | 0.698 | 0.523 | 0.641 | 0.245 |
| vCA3 | Strength | 0.375 | 0.000 | 0.291 | 0.357 |
|  | Eigenvector | 0.403 | 0.000 | 0.266 | 0.399 |
| vCA1 | Strength | 0.581 | 0.028 | 0.906 | 0.693 |
|  | Eigenvector | 0.626 | 0.162 | 0.883 | 0.727 |
| dDCX | Strength | 0.427 | 0.229 | 0.047 | 0.807 |
|  | Eigenvector | 0.317 | 0.297 | 0.031 | 0.783 |
| vDCX | Strength | 0.287 | 0.826 | 0.197 | 0.000 |
|  | Eigenvector | 0.295 | 0.874 | 0.281 | 0.000 |

NC: non-CUS-exposed, CUS: chronic unpredictable stress-exposed, ACC: anterior cingulate cortex, PRL: prelimbic cortex, IL: infralimbic cortex, NAC: nucleus accumbens core, NAS: nucleus accumbens shell, BLA: basolateral amygdala, LA: lateral amygdala, CeA: central amygdala, dDG: dorsal dentate gyrus, dCA3: dorsal cornu ammonis 3, dCA2: dorsal cornu ammonis 2, dCA1: dorsal cornu ammonis 1, dDCX: doublecortin-expressing cells in the dorsal dentate gyrus, vDG: ventral dentate gyrus, vCA3: ventral cornu ammonis 3, vCA1: ventral cornu ammonis 1, vDCX: doublecortin-expressing cells in the ventral dentate gyrus.

***Similarity between networks was low overall, with distinct regions contributing the most to dissimilarity depending on sex and stress***

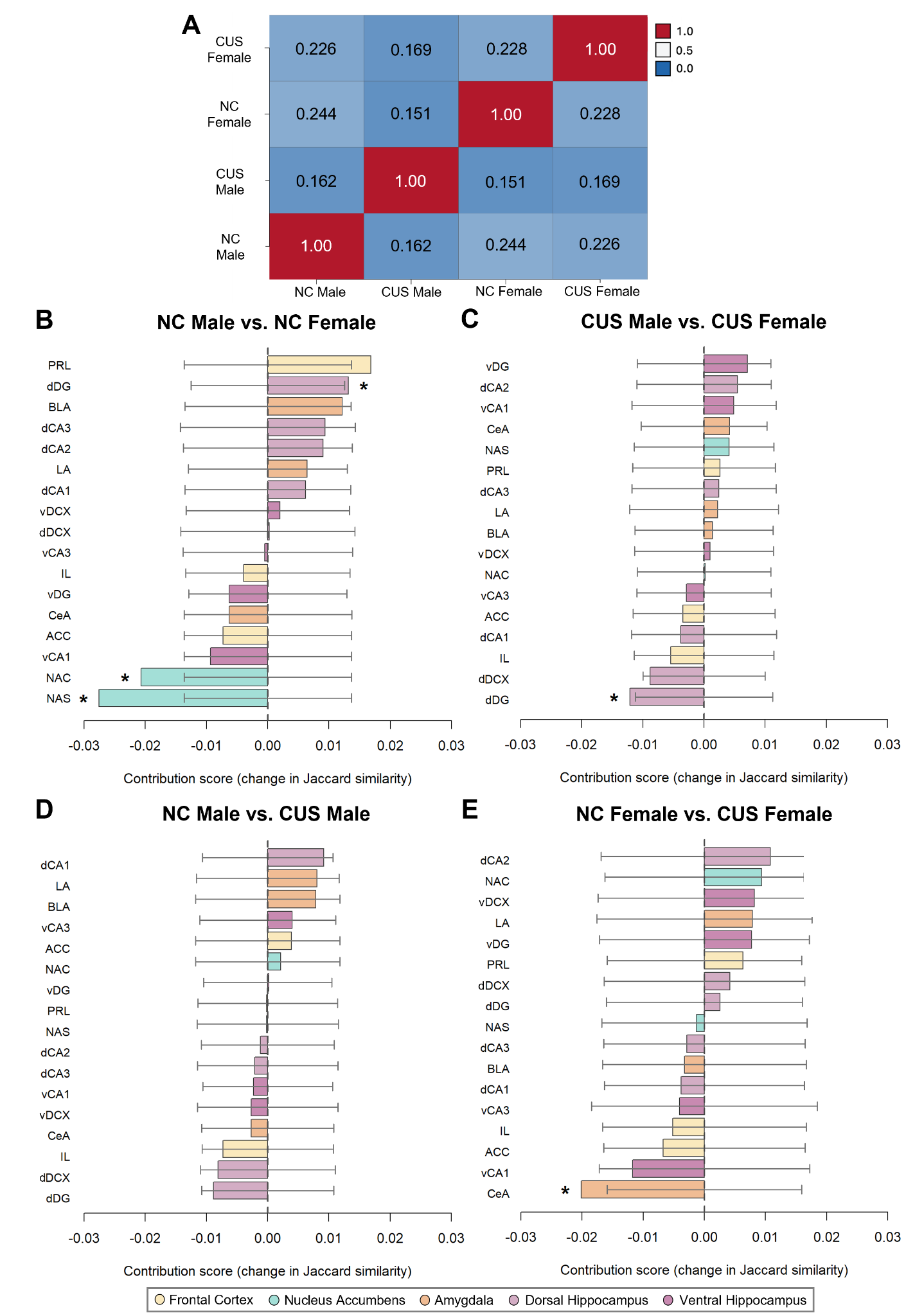

**Supplemental Figure 4. Jaccard network similarity matrix comparing functional networks across groups (A) and regional contribution analysis comparing NC networks across sex (B), CUS networks across sex (C), male networks across CUS (D), and female networks across CUS (E).** In the Jaccard matrix, identical networks have a value of 1.0 and networks with no shared features would have a value of 0.0. Functional networks were largely dissimilar between groups, with the most similar networks being NC females and CUS females, and the least similar networks being NC females and CUS males. In the regional contribution analysis, for a given region, a negative contribution score indicates that that region is contributing to similarity between the networks and a positive contribution score indicates that that region is contributing to difference between the networks. NC: non-CUS-exposed, CUS: chronic unpredictable stress-exposed, ACC: anterior cingulate cortex, PRL: prelimbic cortex, IL: infralimbic cortex, NAC: nucleus accumbens core, NAS: nucleus accumbens shell, BLA: basolateral amygdala, LA: lateral amygdala, CeA: central amygdala, dDG: dorsal dentate gyrus, dCA3: dorsal cornu ammonis 3, dCA2: dorsal cornu ammonis 2, dCA1: dorsal cornu ammonis 1, dDCX: doublecortin-expressing cells in the dorsal dentate gyrus, vDG: ventral dentate gyrus, vCA3: ventral cornu ammonis 3, vCA1: ventral cornu ammonis 1, vDCX: doublecortin-expressing cells in the ventral dentate gyrus. Bars that extend beyond the error bars have a p<0.05, * indicates survival of FDR correction.

We found largely dissimilar networks across sex and stress (Supplemental Fig. 4A, NC females vs. NC males: J=0.244; CUS females vs. CUS males: J=0.169; NC females vs. CUS females: J=0.228; NC males vs. CUS males: J=0.162). The regional contribution analysis comparing the NC networks in response to the ambiguous context revealed that only the nucleus accumbens contributed to similarity between males and females (NAS: p=0.001, NAC p=0.003, Supplemental Fig. 4B). However, the PRL and dDG contributed to dissimilarity (PRL: p=0.018, dDG: 0.044). These data suggest that although the functional networks recruited in response to the ambiguous context are sex-specific even in the absence of stress exposure, the role of the nucleus accumbens is similar between male and female networks. In comparing the networks between males and females after CUS, the dDG significantly contributed to network similarity (p=0.039; Supplemental Fig. 4C), suggesting consistent CUS-induced changes to the dDG’s role across sex.

No regions significantly contributed to similarity or difference between NC and CUS male networks (all p’s>0.092, Supplemental Fig. 4D). The CeA contributed significantly to similarity between NC and CUS female networks (p=0.017, Supplemental Fig. 4E), suggesting that the role of the CeA in female networks remains stable with CUS exposure.

***Regardless of CUS exposure, freezing to the ambiguous context in males was predicted by activity in the amygdala and dentate gyrus in region-specific ways.***

In NC males, activity in the CeA and vDG were each independently significantly positively correlated with freezing (p<0.047). Analysis of region pairs revealed 7 significant correlations between freezing and activity in the CeA and vDG paired with the dCA3, vCA1, or dDCX, all of which were positive except dDCX-vCA3 (all p’s<0.044, Supplemental Table 4). In CUS males, activity in the dDG and LA were independently positively correlated with freezing (p<0.048). Analysis of combinations of two regions revealed 17 significant positive correlations between freezing and activation (all p’s<0.043). The majority of significant correlations (13) included the LA and/or dDG paired with the dCA3 or vDG, among other subregions of the nucleus accumbens, amygdala, frontal cortex, and hippocampus (Supplemental Table 5). However, none of these correlations survived FDR correction.

***Freezing to the ambiguous context in CUS females was driven by activation of the vCA1, but no regions were significantly associated with freezing in NC females.***

In CUS females, activity of vCA1 alone was significantly positively correlated with freezing (p=0.030). Analysis of combinations of two regions revealed significant positive correlations between freezing and 6 region pairs that all include the vCA1 (all p’s<0.048) (Supplemental Table 6). However, no correlations survived FDR correction. No individual regions or region pairs significantly correlated with freezing in NC females (all p’s>0.070).

| **Supplemental Table 4.**  Combinations of regions that best predict freezing in **NC Males.** | | | | | |
| --- | --- | --- | --- | --- | --- |
| # Regions | Regions | R | Adjusted R^2^ | F | p |
| 1 | CeA | 0.6902 | 0.4015 | 6.3667 | 0.0396 |
|  | vDG | 0.6737 | 0.3758 | 5.8162 | 0.0467 |
| 2 | dDCX-CeA | 0.7844 | 0.6577 | 16.3692 | 0.0049 |
|  | vDG-CeA | 0.7208 | 0.4498 | 7.5409 | 0.0287 |
|  | vCA1-CeA | 0.6800 | 0.4265 | 6.9498 | 0.0336 |
|  | dDCX-vDG | 0.6871 | 0.4151 | 6.6787 | 0.0362 |
|  | dCA3-vDG | 0.7774 | 0.3928 | 6.1747 | 0.0419 |
|  | dCA3-CeA | 0.5745 | 0.3890 | 6.0927 | 0.0429 |
|  | dCA3-vCA1 | 0.6645 | 0.3867 | 6.0437 | 0.0436 |

Single regions and region pairs with p<0.05. NC: non-CUS-exposed, CeA: central amygdala, dCA3: dorsal cornu ammonis 3, dDCX: doublecortin-expressing cells in the dorsal dentate gyrus, vDG: ventral dentate gyrus, vCA1: ventral cornu ammonis 1.

| **Supplemental Table 5.**  Combinations of regions that best predict freezing in **CUS Males.** | | | | | |
| --- | --- | --- | --- | --- | --- |
| # Regions | Regions | R | Adjusted R^2^ | F | p |
| 1 | dDG | 0.7275 | 0.4621 | 7.8714 | 0.0263 |
|  | LA | 0.6704 | 0.3708 | 5.7144 | 0.0481 |
| 2 | NAC-LA | 0.7854 | 0.6437 | 15.4553 | 0.0057 |
|  | vDG-LA | 0.7242 | 0.5459 | 10.6158 | 0.0139 |
|  | BLA-LA | 0.7391 | 0.5166 | 9.5497 | 0.0176 |
|  | dDG-IL | 0.7483 | 0.4940 | 8.8096 | 0.0209 |
|  | dDG-LA | 0.7409 | 0.4703 | 8.1024 | 0.0248 |
|  | dCA3-vDG | 0.7416 | 0.4554 | 7.6894 | 0.0276 |
|  | dCA3-dDG | 0.7133 | 0.4458 | 7.4344 | 0.0295 |
|  | IL-LA | 0.7667 | 0.4434 | 7.3732 | 0.0300 |
|  | dCA1-BLA | 0.7828 | 0.4309 | 7.0561 | 0.0326 |
|  | dDG-NAC | 0.6828 | 0.4267 | 6.9538 | 0.0336 |
|  | dCA1-dDG | 0.6839 | 0.4235 | 6.8759 | 0.0343 |
|  | NAS-LA | 0.5728 | 0.4208 | 6.8112 | 0.0349 |
|  | dDG-NAS | 0.6408 | 0.4102 | 6.5650 | 0.0374 |
|  | dDG-vDG | 0.6904 | 0.4025 | 6.3889 | 0.0394 |
|  | dCA3-BLA | 0.7151 | 0.4012 | 6.3596 | 0.0397 |
|  | dDCX-vCA3 | -0.7454 | 0.3975 | 6.2778 | 0.0407 |
|  | dCA3-LA | 0.6710 | 0.3896 | 6.1069 | 0.0428 |

Single regions and region pairs with p<0.05. CUS: chronic unpredictable stress-exposed, IL: infralimbic cortex, NAC: nucleus accumbens core, NAS: nucleus accumbens shell, BLA: basolateral amygdala, LA: lateral amygdala, dDG: dorsal dentate gyrus, dCA3: dorsal cornu ammonis 3, dCA1: dorsal cornu ammonis 1, dDCX: doublecortin-expressing cells in the dorsal dentate gyrus, vDG: ventral dentate gyrus, vCA3: ventral cornu ammonis 3.

| **Supplemental Table 6.**  Combinations of regions that best predict freezing in **CUS Females.** | | | | | |
| --- | --- | --- | --- | --- | --- |
| # Regions | Regions | R | Adjusted R^2^ | F | p |
| 1 | vCA1 | 0.6816 | 0.3976 | 6.9403 | 0.0300 |
| 2 | vCA1-LA | 0.6876 | 0.3563 | 6.5342 | 0.0309 |
|  | vCA1-vCA3 | 0.5508 | 0.3411 | 6.1776 | 0.0347 |
|  | vCA1-CeA | 0.6120 | 0.3221 | 5.7506 | 0.0400 |
|  | dCA2-vCA1 | 0.6748 | 0.3199 | 5.7047 | 0.0407 |
|  | dCA3-vCA1 | 0.6194 | 0.3133 | 5.5629 | 0.0427 |
|  | vCA1-NAS | 0.4607 | 0.2964 | 5.2123 | 0.0483 |

Single regions and region pairs with p<0.05. CUS: chronic unpredictable stress-exposed, NAS: nucleus accumbens shell, LA: lateral amygdala, CeA: central amygdala, dCA3: dorsal cornu ammonis 3, dCA2: dorsal cornu ammonis 2, vCA3: ventral cornu ammonis 3, vCA1: ventral cornu ammonis 1.

| **Supplemental Table 7.**  Region pairs exhibiting significant group differences in linear models. | | | | | |
| --- | --- | --- | --- | --- | --- |
| Region Pair | F | p | Group | R | p |
| CeA-ACC | 3.2685 | **0.0372** | NC Males | 0.5869 | 0.0966 |
|  |  |  | CUS Males | 0.5412 | 0.1321 |
|  |  |  | NC Females | -0.2494 | 0.4871 |
|  |  |  | CUS Females | 0.1881 | 0.5797 |
| vCA1-LA | 3.2823 | **0.0348** | NC Males | 0.1539 | 0.6926 |
|  |  |  | CUS Males | 0.5497 | 0.1252 |
|  |  |  | NC Females | -0.0519 | 0.8867 |
|  |  |  | CUS Females | 0.6486 | **0.0309** |
| vCA1-ACC | 3.5503 | **0.0287** | NC Males | 0.3980 | 0.2887 |
|  |  |  | CUS Males | 0.1283 | 0.7421 |
|  |  |  | NC Females | -0.0533 | 0.8837 |
|  |  |  | CUS Females | 0.3754 | 0.2553 |
| vCA1-dDCX | 3.9461 | **0.0174** | NC Males | 0.5053 | 0.1653 |
|  |  |  | CUS Males | -0.4670 | 0.2050 |
|  |  |  | NC Females | 0.3641 | 0.3010 |
|  |  |  | CUS Females | 0.5512 | 0.0788 |

Statistics for region pairs exhibiting interactions with p<0.05 and linear modeling results by group for each significant region pair. Significant p-values are bolded. NC: non-CUS-exposed, CUS: chronic unpredictable stress-exposed, ACC: anterior cingulate cortex, LA: lateral amygdala, CeA: central amygdala, dDCX: doublecortin-expressing cells in the dorsal dentate gyrus, vCA1: ventral cornu ammonis 1.

***Linear models exhibit group differences regardless of statistical significance of individual functions***

Four linear models exhibited significant group differences: CeA-ACC, vCA1-LA, vCA1-ACC, and vCA1-dDCX (Fig. 7, Supplemental Table 7). Activation of these region pairs did not significantly correlate with freezing within individual groups, with the exception of vCA1-LA in CUS females (p=0.031).
